# Tonic resting-state hubness supports high-frequency activity defined verbal-memory encoding network in epilepsy

**DOI:** 10.1101/660696

**Authors:** Chaitanya Ganne, Walter Hinds, James Kragel, Xiaosong He, Noah Sideman, Youssef Ezzyat, Michael R Sperling, Ashwini Sharan, Joseph I Tracy

**Author notes:** Corresponding Author Joseph I. Tracy, Ph.D., Professor, Departments of Neurology and Radiology, Director, Clinical Brain Mapping and Cognitive Neuroscience, Laboratory, Thomas Jefferson University, Sidney Kimmel Medical College, 901 Walnut Street, Suite 447, Philadelphia, PA, 19107.

## Abstract

High-frequency gamma activity of verbal-memory encoding using invasive-electroencephalogram coupled has laid the foundation for numerous studies testing the integrity of memory in diseased populations. Yet, the functional connectivity characteristics of networks subserving these HFA-memory linkages remains uncertain. By integrating this electrophysiological biomarker of memory encoding from IEEG with resting-state BOLD fluctuations, we estimated the segregation and hubness of HFA-memory regions in drug-resistant epilepsy patients and matched healthy controls. HFA-memory regions express distinctly different hubness compared to neighboring regions in health and in epilepsy, and this hubness was more relevant than segregation in predicting verbal memory encoding. The HFA-memory network comprised regions from both the cognitive control and primary processing networks, validating that effective verbal-memory encoding requires multiple functions, and is not dominated by a central cognitive core. Our results demonstrate a tonic intrinsic set of functional connectivity, which provides the necessary conditions for effective, phasic, task-dependent memory encoding.

**Highlights:**

1. High frequency memory activity in IEEG corresponds to specific BOLD changes in resting-state data.
2. HFA-memory regions had lower hubness relative to control brain nodes in both epilepsy patients and healthy controls.
3. HFA-memory network displayed hubness and participation (interaction) values distinct from other cognitive networks.
4. HFA-memory network shared regional membership and interacted with other cognitive networks for successful memory encoding.
5. HFA-memory network hubness predicted both concurrent task (phasic) and baseline (tonic) verbal-memory encoding success.

## Introduction

Successful episodic verbal memory encoding, termed the ‘Subsequent Memory Effect’ (SME) is associated with high-frequency gamma activity (ranging from 30-100Hz) captured using power spectral and synchronization studies during invasive electroencephalography (IEEG) in association with neuronal firing (Burke JF et al., 2015;Kucewicz MT et al., 2017;Lega BC et al., 2012;Long NM et al., 2014;Sederberg PB et al., 2006;Serruya MD et al., 2014;Solomon EA et al., 2017;Solomon EA et al., 2019). Despite these well-established linkages, relatively little is known about the whole-brain organization of HFA regions and the degree to which these regions possess either distinct network features relative to other brain areas, or rely on abnormal organizational properties due to epilepsy. Kucewicz and colleagues demonstrated that memory related broadband gamma activity centroids were in the range of 30-100Hz (Kucewicz MT, et al., 2017). Two recent studies on human memory network analysis with a pooled cohort of patients similar to that included in this study, have shown that successful memory encoding is associated with asynchronous organization of the memory network in gamma activity less than 100Hz and synchronous theta activity, with widespread high gamma activity considered to be the hallmark of memory encoding (Solomon EA, et al., 2017;Solomon EA, et al., 2019). In this project, we were interested in investigating the intrinsic BOLD correlations of a high frequency network already known to play a major role in human memory, all toward determining if such networks possess distinctive whole-brain organization properties.

While evidence exists to suggest that memory-relevant regions in the epileptic brain can take up normative network roles in an effort to preserve function (Jin SH et al., 2015;Powell HW et al., 2007;Solomon EA, et al., 2017;Tracy J et al., 2014), the network properties characterizing these memory regions in health and in disease are unclear. In this study, we integrate IEEG with resting-state fMRI (rsfMRI) data to characterize the network architecture of a task-defined HFA-memory network and understand its relationship to the functional connectivity of a broader set of intrinsic resting-state networks. Graph theory has been a useful tool for mapping functional networks and characterizing their properties during task and at rest. Three important graph indices that are essential in characterizing brain regions and their functionality include: (1) clustering coefficient (CC), which captures network segregation and local information processing and (2) betweenness centrality (BC), which captures hubness and the degree of importance held by a region that connects two or more modules and (3) participation coefficient (PC) which measures the extent to which regions within a network connect to networks other than its own, with a higher participation coefficient indicating that the regions connect with a variety of other networks. While BC indicates regions that have considerable influence within a network by virtue of their control over information passing between others, it does not reveal anything about the diversity of inter-network connections of individual nodes. In contrast, PC helps identify influential nodes in a network that are likely to be highly connected to other networks and, as a result, communicate and exert influence over these other networks. In the setting of our paper, PC helps verify the actual level of interaction of the HFA network with the intrinsic networks (Power JD et al., 2013). Through network neuroscience measures of segregation, hubness and interaction, we define and broaden our understanding of the network characteristics that support memory encoding and reveal the contribution of multiple intrinsic networks to successful memory encoding.

Prior studies combining IEEG and fMRI have shown a good correspondence between HFA and blood oxygen level dependent (BOLD) contrast signal, offering a key link between electrophysiological and hemodynamic properties of human cognition (Axmacher N et al., 2008;Jacques C et al., 2016;Khursheed F et al., 2011;Logothetis NK et al., 2001;Rugg MD et al., 2002). Integrating IEEG with rsfMRI has the advantage of sampling the wider brain regions, allowing us to test the correspondence between the information embedded in the faster dynamics of IEEG data (i.e., HFA) and the slower BOLD response (Mizuhara H, 2012;Mizuhara H et al., 2005). In this study, we address four specific questions: First, how does the resting-state functional connectivity (rsFC) of the HFA-memory network involved in verbal-memory encoding differ from neighboring regions that lack significant SME? Second, are the network characteristics of HFA-memory regions in epilepsy patients generalizable, or do they differ from those of matched healthy controls? Third, by discovering the key features of the HFA-memory network and studying their relationship to other intrinsic networks, can we reveal the trait-like properties of the intrinsic state, as well as the constituent cognitive processes that are necessary for effective memory encoding? Fourth, are HFA-memory network characteristics associated with and able to predict clinically relevant verbal-memory performance in epilepsy?

## METHODS

### Participants

Thirty-seven patients with drug-resistant epilepsy were recruited from the Thomas Jefferson University Comprehensive Epilepsy Center. They underwent IEEG implantation (subdural, depth, or both) to localize the epileptogenic zone and guide potential surgical management of their seizures (Table 1). Sites of implantation were determined by multimodal pre-surgical evaluation including neurological history and examination, video-EEG, MRI, PET, and neuropsychological testing (Sperling MR et al., 1996). Specific to this study, patients underwent a 3T-MRI scan involving resting state (rsfMRI), a high resolution anatomical image (T1MPRAGE) (Philips Achieva, Amsterdam, Netherlands), pre-surgical neuropsychological assessment (NPA) to provide a baseline indication of the patients verbal-memory skills, followed by IEEG implantation and monitoring (Nihon Kohden EEG-1200, Irvine, CA) during which patients participated in verbal episodic memory testing (free-recall paradigm). This provided behavioral measures at two stages of clinical evaluation: percent recall from a word-list memory test administered simultaneously with IEEG recording (P.REC), and the sum-total of words recalled from a similar memory test administered during baseline neuropsychological testing (CVLT Total Learning - CVLT-TL) (Supplementary Methods, Data Acquisition).

Patients were excluded from the study for any of the following reasons: previous brain surgery; medical illness with central nervous system impact other than epilepsy; contraindications to MRI; or hospitalization for major psychiatric disorders as per Diagnostic and Statistical Manual of Mental Disorders-V, except depressive disorders were allowed given the high comorbidity of depression and epilepsy (Tracy JI et al., 2007). Healthy controls (HC, N=37) were recruited to match the patients in age, gender, handedness, and education. Since IEEG data can only be acquired from patients with epilepsy undergoing implantation, we needed to test if the network characteristics of HFA-memory regions in epilepsy patients are generalizable (i.e., do they differ from those of matched, healthy controls?). Hence, regions homologous to the HFA-memory network and the neighboring controls nodes (described later) were identified in the healthy controls and their network characteristics were tested and compared to patients. All participants provided written informed consent as per the TJU Institutional Review Board requirements.

### IEEG acquisition and free-recall testing

IEEG data [subdural grids and strips (10mm resolution) or depth electrodes (5-10mm resolution)] were recorded from the patients performing delayed free recall of categorized and unrelated word lists in three distinct phases of encoding, delay, and retrieval (https://github.com/pennmem/wordpool). The task was presented using PyEPL software and the data were simultaneously recorded using the Nihon-Kohden IEEG system [sampling rate >500Hz] (Geller AS et al., 2007). Participants were instructed to commit each list of words to memory. Recalled responses were digitally recorded and parsed/scored offline using the University of Pennsylvania Total Recall program (http://memory.psych.upenn.edu/TotalRecall). Participants performed up to 25 recall trials in a single recall session. During encoding, lists of 12 words were presented (each word for 1600ms followed by a blank inter-stimulus interval of 750 to 1000ms). Following a math distractor task, the participant recalled as many words as possible from the most recent list presented (recall period-30s). (Figure 1). A measure of immediate recall, representing the percentage of words recalled across the multiple trials was utilized (percent recall, P.REC).

**Figure 1:**
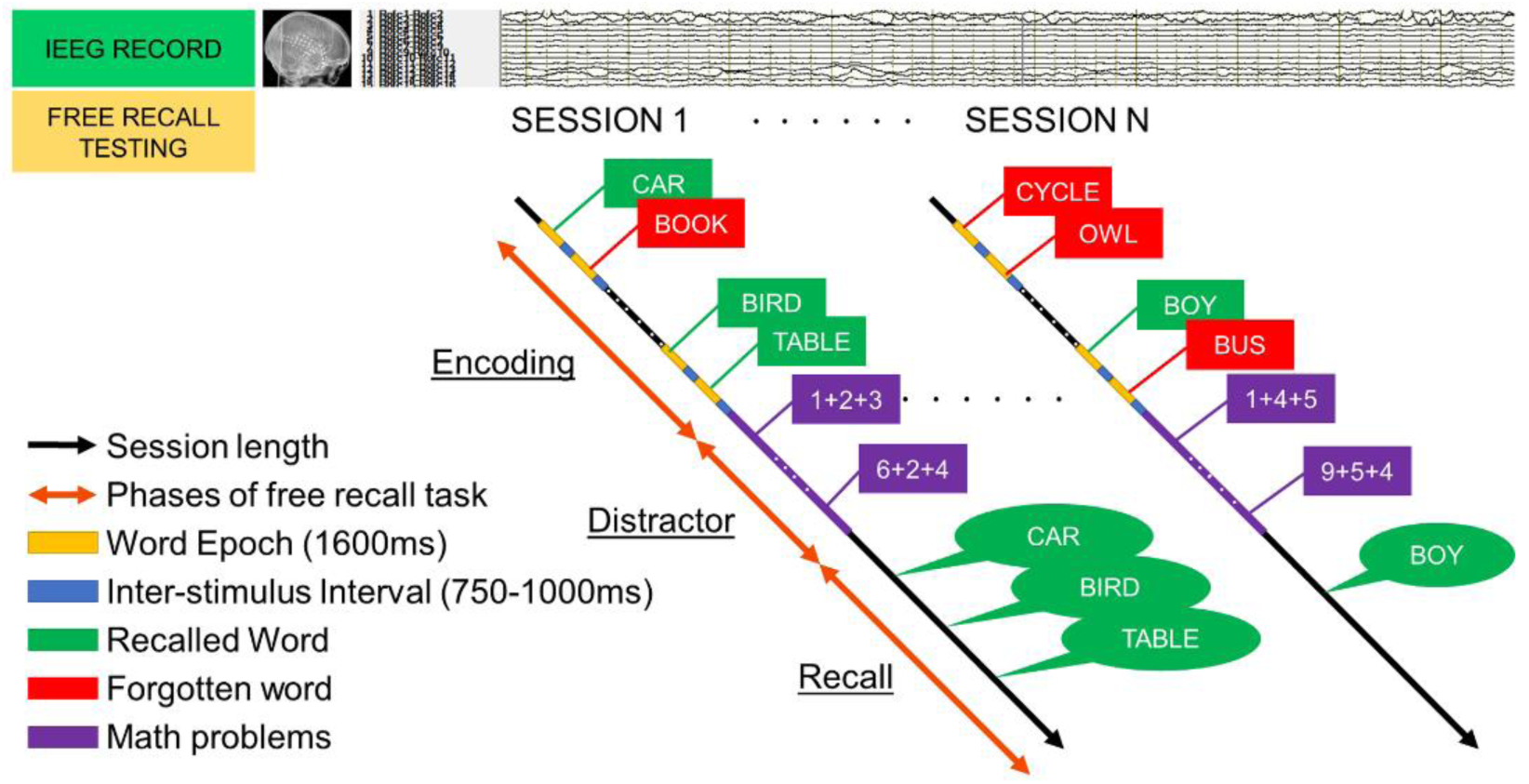
Free-recall task design. This is a schematic representation of simultaneous IEEG acquisition and encoding/free recall testing sessions administered to the epilepsy patients in the EMU. Time-synced markers for stimulus presentation were recorded on the IEEG signal. There were 3 phases of the task: 1) encoding, where a list of new random words is presented to the subject and they were asked to commit them to memory. During encoding, lists of 12 words were presented (each word for 1600ms followed by a blank inter-stimulus interval (ISI) of 750 to 1000ms). 2) distractor task, involving a 20s post-encoding delay period containing an arithmetic task was given to disrupt memory for end-of-list items (patients solved math problems of the form A+B+C=X using a response keypad). 3) recall, where the subject was asked to recall the words presented during the encoding phase. This design provided information about IEEG activity during the words that were successfully encoded verses those that were forgotten. Recalled responses were digitally recorded and parsed/scored offline using the University of Pennsylvania Total Recall program (http://memory.psych.upenn.edu/TotalRecall). A measure of immediate recall, representing the percentage of words recalled across the multiple trials was estimated (percent recall, P.REC). Participants performed up to 25 recall trials (i.e., lists) in a single recall session.

Signals recorded were first referenced to a common contact placed intracranially, on the scalp, or mastoid process. To eliminate potentially confounding large-scale artifacts and noise on the reference channel and to estimate the power analyses, we calculated the bipolar derivative where each electrode was linked to the next along a chain. Electrodes which were a part of the epileptogenic zone were excluded, so as not to confound power spectral changes related to memory with those caused by epileptogenic activity (ictal or interictal). Data was downsampled to 500Hz uniformly. The bipolar time-series derived from the implanted electrodes were band-pass filtered with a 4th order Butterworth band-reject filter from 58 to 62Hz to exclude line-noise. To quantify memory-related changes in spectral power, complex-valued Morlet wavelets (wave number 6) were convolved with the down-sampled (200Hz) bipolar IEEG signals to obtain magnitude information. Ten wavelets with center frequencies logarithmically-spaced between 44 and 100Hz were used to estimate changes in HFA power. The average spectral power for each encoding event was computed from the onset of each stimulus for a duration of 1600ms, with a buffer of 1000ms. Prior to computing the subsequent memory effect (SME), power values for each bipolar pair were log-transformed and normalized to have zero mean and unit standard deviation within each session. A paired t-test was performed between the HFA power of words that were remembered and words that were forgotten at each bipolar pair. A significant increase in the HFA power provided a measure of the subsequent memory effect (SME).

Subsequently, anatomic localization of bipolar pairs (computed as the mid-point between the two contacts) was accomplished using 2 independent processing pipelines for depth and surface electrodes which were later transformed to MNI space similar to previous studies (Burke JF et al., 2013;Kragel JE et al., 2017).

### Functional MRI data acquisition and preprocessing

All participants (patients and healthy controls) underwent a rsfMRI scan using a single-shot echo-planar gradient echo sequence (120 volumes; 34 slices; TR = 2.5s, TE = 35ms; flip angle (FA) = 90°, FOV=256 mm, 128×128 matrix, resolution: 2×2×4mm) in a 3T MRI scanner (Philips Achieva). T1-MPRAGE images (180 slices, 256×256 isotropic 1mm voxels; TR/TE/FA=640ms/3.2ms/8°, FOV=256 mm) were also collected. Patients were instructed to stay awake, keep their eyes closed, and stay relaxed. Patients who reported seizures within 24 hours of the scheduled MRI appointment were not scanned. The scans were performed mid-day so that patients were cooperative, alert, and not fatigued. The resting-state scan was performed immediately after the T1 image, within 15 minutes of starting the scan, to help ensure that there was no fatigue from being in the scanner. Subject’s promptness in responding to oral commands immediately after the resting-state scan was used to confirm the participant’s level of alertness. If there was any indication that the participant fell asleep or was not awake and alert, their data were excluded from analyses.

All imaging data were preprocessed using Data Processing Assistant for rsfMRI Advanced Edition (http://www.rfmri.org/DPARSF)(Yan CG et al., 2010), a MATLAB toolbox based on Statistical Parametric Mapping-8 (http://www.fil.ion.ucl.ac.uk/spm/software/spm8) in the following order: removal of first three fMRI volumes to maintain BOLD signal homogeneity, slice timing correction in an interleaved manner with the mid-time slice as a reference, realignment and co-registration of T1-MPRAGE and the corresponding rsfMRI, brain extraction using Brain Extraction Toolkit (https://github.com/neurolabusc/MRIcro), segmentation of the T1MPRAGE with neo-segmentation algorithm+DARTEL (http://www.fil.ion.ucl.ac.uk/spm/software/spm8), nuisance covariate regression using Friston’s 24-parameter model (Friston KJ et al., 1996), cerebral spinal fluid and white matter mask regression using CompCor (Ashburner J et al., 2005;Yan CG et al., 2013), filtering resultant fMRI data between 0.008-0.1Hz, normalization to the MNI space and smoothing using a Gaussian Kernel of 4mm full width half maximum (FWHM).

### Graph Theory analysis

Using the Lausanne’s atlas of 234 cortical and subcortical regions of interest (ROI), a 234×234 correlation-matrix was calculated at individual subject level, which was subsequently used to construct weighted undirected graph. Minimum Spanning Tree (MST) based networks were derived to ensure the same number of connected nodes for all subjects, allowing for reliable group-level comparisons and yielding a series of graphs with connection density ranging from 5% to 50% in increments of 1% (Stam C et al., 2014;Tewarie P et al., 2015;van Diessen E et al., 2013). The density range of 5% to 50% was chosen for the following reasons: (1) network measures are relatively constant over this range (Alexander-Bloch AF et al., 2010); (2) previous work has suggested that above a density of 50% graphs become more random (Humphries MD et al., 2006) and prone to non-biological features and noise (Kaiser M et al., 2006). Using these graphs, CC, BC and PC were calculated using Brain Connectivity Toolbox (www.brain-connectivity-toolbox.net) (Rubinov M et al., 2010).

### Defining HFA-memory regions (MEM) and their neighboring control nodes (CNs) for rsfMRI analysis

Brain regions where IEEG showed significant SME with increased HFA were termed HFA-memory regions (MEM). Brain regions where IEEG recorded HFA, but the results did not survive statistical significance for successful memory encoding were termed Control Nodes (CNs).

The 3731 IEEG bipolar contacts for all the patients were superimposed into a single MNI coordinate space, referred to as the “super-brain” (FIGURE 2A and B).

**Figure 2:**
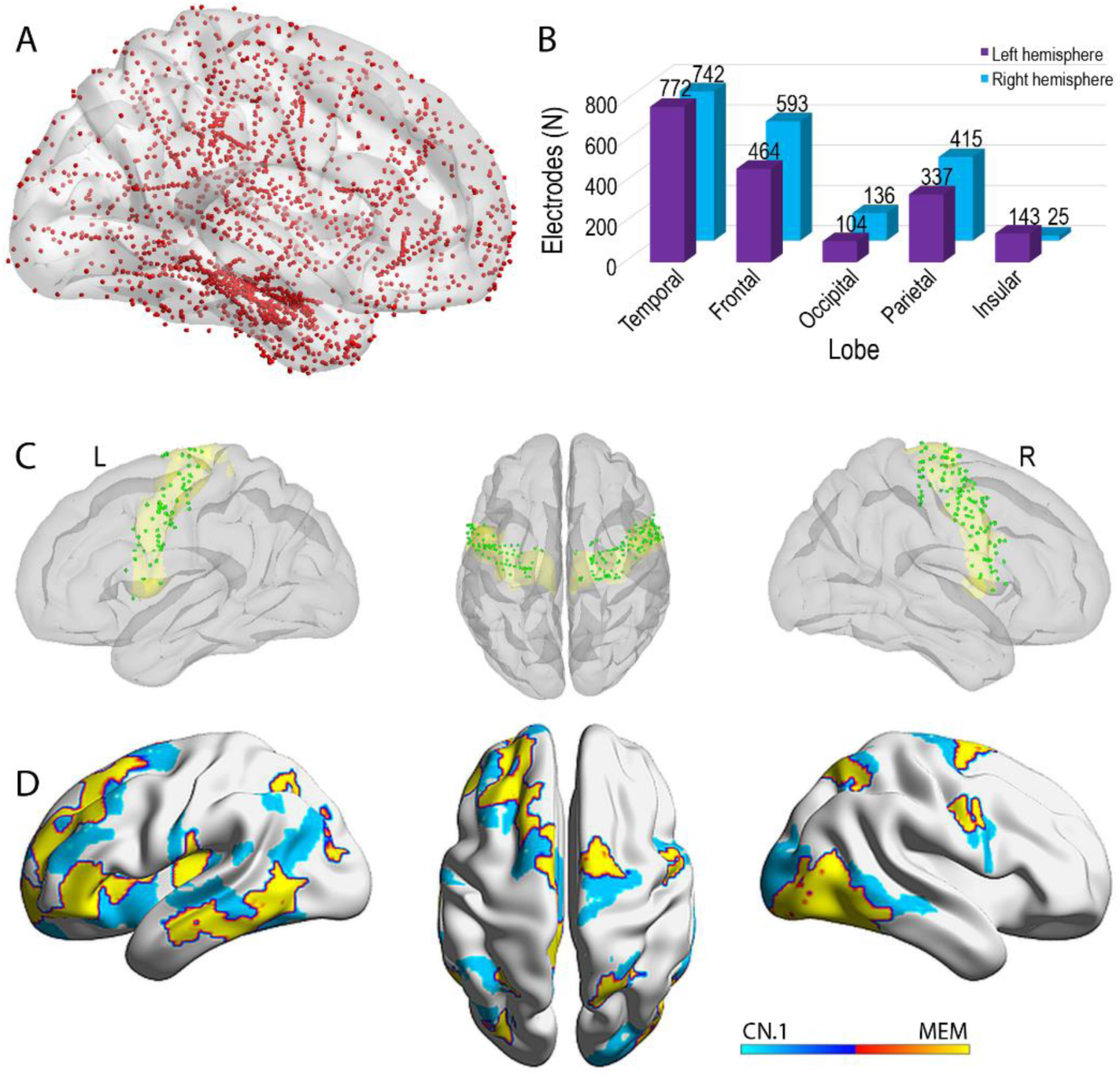
IEEG implantation and electrode assignment to brain parcels. (A) The 3731 IEEG bipolar contacts for all the patients were superimposed into MNI coordinate space, referred to as the “super-brain”. (B) Shows the distribution of the IEEG bipolar contacts in the different lobes across all subjects (N=37). (C) Is an example of the assignment of the bipolar IEEG contacts to the nearest parcel of the Lausanne’s atlas. Here we note a total of 188 electrodes (green spheres) from 37 patients were assigned to bilateral precentral gyri (yellow regions) of the super-brain. The minimum number of bipolar contacts in any given parcel were no less than 16 in number. (D) Shows the HFA-memory regions (MEM) and their corresponding Control Nodes (CNs). The warm yellow color represents the HFA-memory regions (MEM) and the cool cyan color represents the closest set of controls nodes (CN.1). The brain regions were illustrated in BrainNet Viewer (http://www.nitrc.org/projects/bnv/).

The MNI coordinates of these contacts were then warped to the nearest 3D Cartesian coordinates in the Euclidean space of each of the ROIs of the Lausanne atlas (Hagmann P et al., 2008) (FIGURE 2C). In order to avoid the probability of the algorithm wrongly assigning the electrodes to subcortical structures, we excluded these regions, yielding an atlas with 222 implant-relevant regions. Each electrode coordinate was warped to the coordinate of the nearest ROI of the Lausanne atlas. This allowed us to group the 3731 contacts into their respective 222 ROIs. Once this was done, the t-values of the multiple electrodes within each ROI were submitted to a positive tailed one-sample t-test to determine if there was a significant increase in HFA power between words that were encoded and words that were forgotten. To account for multiple comparisons, the P-values generated for the 222 ROIs were subjected to FDR correction (Genovese CR et al., 2002). The FDR corrected significant ROIs were considered to exhibit memory-relevant encoding effects, henceforth referred to as MEM ROIs.

For every MEM, we identified the nearest 5 CNs based on their Euclidian distance, as these neighboring CNs had the highest chance of being implanted and providing contrasting HFA recordings. The ROIs closest to MEM were grouped as first control nodes (CN.1) and second closest as CN.2. To avoid the bias of the distance from the MEM, we chose a random ROI among the first 5 CNs, and refer to these as the ‘random distance control nodes’ (CN.RAND).

### Hubness and Participation coefficient of the HFA-memory network in comparison with the intrinsic networks

Since the HFA-memory network was found to have regions distributed widely over the brain, we were interested in looking at whether the HFA-memory regions had membership in other intrinsic networks. Hence, we assigned the different regions of the Lausanne’s atlas to the different intrinsic networks as defined by Gu et al (Kauffmann L et al., 2015) and estimated the percentage contribution of the different intrinsic networks to the HFA-memory network (i.e., percentages avoid bias from the different number of ROI’s that constitute each intrinsic network). The networks included cognitive control systems (temporal (TN), frontoparietal (FPN), dorsal attention (DAN), and cinguloopercular (COpN) networks), primary processing networks (sensorimotor (SMN), visual integrated (VIN) and auditory (AN) networks and a task-negative network, the default mode network (DMN).

Noting that the HFA-memory regions do not constitute an intrinsic network, nor a surrogate for any one of them, we estimated the average BC and PC of each of the networks (AvgBC and AvgPC respectively), and sought to determine if the BC and PC of the HFA-memory network differed from that of the intrinsic networks using ANOVA. We ranked the networks according to their AvgBC values. To further understand the nature of the overlap between intrinsic networks and the HFA-memory network, we tested whether the BC of ROIs of an intrinsic network that contributed to the HFA-memory regions differed from those that did not form a part of the HFA memory network.

### Statistical analysis

Chi-square or t-tests, as appropriate, were used to compare the groups on demographic and clinical variables (p<0.05 was considered significant). To assess group (patient vs. controls), region (MEM vs. CN), and group-by-region effects, we ran a two-way ANOVA on the graph indices (CC and BC) at two levels: 1) the ‘composite network’ level and 2) ‘individual region’ level. At a ‘composite network’ level, the average-CC (AvgCC) and average-BC (AvgBC) across the separate MEM and CNs were calculated and served as the dependent variables in the ANOVAs. The ANOVAs contrasting the MEM with the three available CN ROI’s (CN.1, CN.2, and CN.RAND) were run in separate models. At the ‘individual region’ level, the NodalCC and NodalBC of MEM and CNs served as the dependent variables in the ANOVAs. A MEM was considered significantly different from the CNs only when each MEM was shown to be reliably different from all three CNs (CN.1, CN.2, and CN.RAND) using Boolean conjunction analysis.

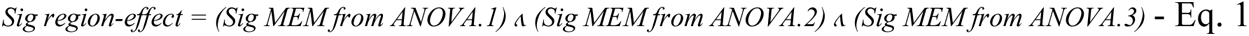

Where *Sig region-effect* is the significant region-effect, ANOVA.1 is tested between MEM and CN.1, ANOVA.2 is tested between MEM and CN.2 and ANOVA.3 is tested between MEM and CN.RAND and Λ is the commutative conjunction operator. This was a stringent requirement for significance ensuring that MEM ROIs with a significant difference in graph indices to only a single CN were not considered valid. Throughout the analysis, multiple comparison correction was calculated using the false discovery rate (pFDR). The difference in AvgBC and AvgPC of the HFA-memory network in comparison with the intrinsic networks was tested using repeated-measures ANOVA with Fisher’s Least Significant Difference (LSD).

### Multivariate machine learning to predict free-recall performance

We used a ‘Random Forest’ (RF) model to determine the relationship between fMRI graph indices (NodalBC and NodalCC, as the predictor variables) and free-recall performance (P.REC and CVLT-TL, tested separately, i.e., the response variable). CVLT-TL is sensitive to memory encoding and immediate recall, comparable in this sense to the P.REC measure. RF is an efficient, supervised machine learning method that identifies both linear and non-linear brain-behavior relationships, allows simultaneous testing of multivariate interactions, and shows superior resilience to over-fitting (Breiman L, 2001). The algorithm exploits random decision trees, which use a subset of the observations through bootstrapping techniques. In short, from the original data set a random selection of the training data is sampled and used to build a model. The model is then tested on the data not included in the training sample, referred to as “out-of-bag”. The random forest algorithm estimates the variable importance (VI) by looking at how much prediction error increases when out-of-bag data for that variable is permuted while all others are left unchanged. (Liaw A et al., 2002). Briefly, we estimated the true VI for each predictor for 1000 repetitions. Next, we established a null distribution of VIs for every predictor by estimating their VI from models trained with the response variable randomly permuted a 1000 times. Based on the true VI, the VI of each predictor from each random permutation, a probability can be estimated based on its corresponding normal cumulative null distribution, with a variable considered significant if the probability is larger than 95%. Lastly, we selected predictors whose VIs were found significant at least 950 times over the 1000 repetitions, to build RF models and predict the response. The association between the predictions and actual scores was tested using a right-tailed Pearson correlation (He X et al., 2018). In addition, stepwise linear regression models on the two response variables were run with the selected predictors from the RF models serving as independent variables, allowing us to explore any directional or ranking effects among the predictors.

## RESULTS

### Demographic and clinical characteristics of the groups (patients and controls)

The demographic details of the patients and healthy controls are listed in Table 1 (also Appendix A). They were matched for age, gender, handedness, years of education, and head micro-movement during rsfMRI (p>0.05) and all participants were awake and alert at the end of the resting state scan and promptly responded to oral commands. Relative to same-age peers, the patients had CVLT-TL performance in the average range (t=48, age-normed), indicating intact memory encoding ability.

**Table 1:** Demographic and clinical characteristics. Both patients and healthy controls were matched for age, gender, handedness, education and head motion. Further demographic data specific to the patients have been enumerated in this table. (M: Mean, SD: Standard Deviation, FOA: Focal Onset Aware Seizures, FOIA: Focal Onset seizures with Impaired Awareness; BTCS: focal seizures progressing to bilateral tonic-clonic seizures, VGNC: Voltage-gated sodium channel blockers: CBZ: carbamazepine, OXC: oxcarbazepine, PHY: phenytoin, GABAa Agonist: Gamma aminobutyric acid a receptor agonist: PB: barbiturates; BZDs: benzodiazepines (diazepam, lorazepam, clonazepam, clobazam); SV2a Receptor-Mediated AEDs: LVA: levetiracetam; CRMP2 Receptor-Mediated AEDs: LCM: lacosamide; VGCC: Voltage-gated calcium channel: PGB: pregabalin; GBP: gabapentin; Multi-action AEDs: VPA: valproate; TPM: topiramate; Pr: Primidone).

| Sample Group<br>(N) | | Patients<br>37 | Controls<br>37 | F/T/$\chi^2$ | P |
| --- | --- | --- | --- | --- | --- |
| Age (M±SD) | | 36.86±10.69 | 33.91±12.65 | 1.08 | 0.28 |
| Gender (M/F) |  | 21/16 | 24/13 | 0.23 | 0.63 |
| Education (years, M±SD) | | 13.67±2.62 | 14.2±1.72 | -1.39 | 0.16 |
| Handedness (R:L) |  | 30/7 | 26/11 | 0.66 | 0.41 |
| Head Motion (FD_Jenkinson) | | 0.12±0.01 | 0.10±0.04 | -0.90 | 0.36 |
| Free-recall Measures |  |  |  |  |  |
| P.REC | | 21.15±9.2% | | | |
| CVLT-TL | | 48.4±11.2% | | | |
| Age at Epilepsy Onset<br>(M±SD; years) | | 20.94±13.19 | - | - | - |
| Duration of Epilepsy (M±SD;<br>years) | | 15.92±10.25 | - | - | - |
| <u>Seizure Type</u> |  |  |  |  |  |
| FOIA |  | 6 | - | - | - |
| FOIA+BTCS |  | 17 | - | - | - |
| FOIA/FOA |  | 4 | - | - | - |
| FOIA/FOA+BTCS |  | 6 | - | - | - |
| FOA+BTCS |  | 4 | - | - | - |
| <u>Anti-Epileptic Drugs</u> |  |  |  |  |  |
| VGNC | CBZ,OXC,LTG,<br>PHT | 24 | - | - | - |
| GABAa Agonist | PB, BZD, Pr | 7 | - | - | - |
| SV2a Receptor Mediated | LVA | 15 | - | - | - |
| CRMP2 Receptor Mediated | LCM | 8 | - | - | - |
| Multi-Action | VPA, TPM,<br>ZNS | 17 | - | - | - |
| VGCC | PGB, GBP | 4 | - | - | - |

### Defining HFA-memory regions and their neighboring control nodes

Brain regions where IEEG showed significant SME with increased HFA were termed HFA-memory regions (MEM). MEM regions included regions of lateral occipital cortex and the inferotemporal surface consisting of the fusiform gyri along with regions of the lateral temporal cortex, dorsolateral prefrontal cortex, precuneus, and cinguloopercular regions. MEM areas also included discrete regions of bilateral lateral neocortices involving the left rostral middle frontal, inferior temporal, cingulate, postcentral, inferior parietal, insular (LrMFG, LITG, LPostCinG, LIstCinG, LIPL, LInsG), bilateral superior frontal, parietal, inferior temporal, and fusiform gyri (B/LSFG, B/LSPG, B/LITG, B/LFusG) (pFDR<0.0058). The corresponding control regions were identified (Table 2 and Figure 2D).

**Table 2:** HFA-memory regions (MEM) and their corresponding control nodes (CN) The 3731 electrodes after being assigned to the different brain regions were tested for significant subsequent memory effect (MEM). Once these regions were derived based on nearest Euclidean distances we determined the three neighboring control nodes (CN.1, CN.2, and CN.RAND). (ROI# - The ROI numbering as per the atlas; MEM – IEEG defined ROIs which were significant for HFA-memory activity; CN – Control nodes that are measured at 3 different Euclidean Distances; R.superiorfrontal (RSFG), R.precentral (RPreCG), R.superiorparietal (RSPG), R.lateraloccipital (RLatOG), R.fusiform (RFusG), R.inferiortemporal (RITG), L.parsopercularis (LparsOp), L.parstriangularis (LparsTri), L.rostralmiddlefrontal (LrMFG), L.superiorfrontal (LSFG), L.posteriorcingulate (LPostCinG), L.isthmuscingulate (LIstCinG), L.postcentral (LPostCG), L.superiorparietal (LSPG), L.inferiorparietal (LIPG), L.fusiform (LFusG), L.inferiortemporal (LITG), L.middletemporal (LMTG), L.insula (LIns), R.lingual (RLinG), R.entorhinal (REntRG), R.middletemporal (RMTG), L.lateralorbitofrontal (LLatOrbFG), L.caudalmiddlefrontal (LcMFG), L.precuneus (LPC), L.entorhinal (LEntRG), L.superiortemporal (LSTG), L.transversetemporal (LTransTG), R.caudalmiddlefrontal (RcMFG), R.isthmuscingulate (RIstCinG), R.inferiorparietal (RIPG), R.parahippocampal (RPHG), L.precentral (LPreCG), L.paracentral (LParaCG), L.supramarginal (LSMG), L.lateraloccipital (LLatOG), L.parahippocampal (LPHG), R.paracentral (RParaCG), R.postcentral (RPostCG), R.pericalcarine (RPeriCaL), R.amygdala (RAmy), L.accumbensarea (LAccu), L.amygdala (LAmy))

| ROI# | MEM | ROI# | CN.1 | ROI# | CN.2 | ROI# | CN.RAND |
| --- | --- | --- | --- | --- | --- | --- | --- |
| 26 | RSFG.7 | 32 | RPreCG.2 | 25 | RSFG.6 | 38 | RParaCG.2 |
| 33 | RPreCG.3 | 36 | RPreCG.6 | 27 | RSFG.8 | 47 | RPostCG.3 |
| 57 | RSPG.4 | 56 | RSPG.3 | 29 | RcMFG.2 | 55 | RSPG.2 |
| 78 | RLatOG.3 | 76 | RLatOG.1 | 44 | RIstCinG.1 | 65 | RIPG.5 |
| 79 | RLatOG.4 | 77 | RLatOG.2 | 58 | RSPG.5 | 74 | RPeriCaL1 |
| 80 | RLatOG.5 | 83 | RLinG.3 | 65 | RIPG.5 | 77 | RLatOG.2 |
| 84 | RFusG.1 | 86 | RFusG.3 | 66 | RIPG.6 | 81 | RLinG.1 |
| 85 | RFusG.2 | 89 | REntRG.1 | 81 | RLinG.1 | 93 | RITG.3 |
| 87 | RFusG.4 | 95 | RMTG.1 | 82 | RLinG.2 | 94 | RITG.4 |
| 94 | RITG.4 | 96 | RMTG.2 | 88 | RPHG.1 | 115 | RAmy |
| 120 | LparsOrb.1 | 117 | LLatOrbFG.2 | 93 | RITG.3 | 116 | LLatOrbFG.1 |
| 124 | LparsTri.1 | 119 | LLatOrbFG.4 | 97 | RMTG.3 | 129 | LrMFG.3 |
| 128 | LrMFG.2 | 127 | LrMFG.1 | 118 | LLatOrbFG.3 | 134 | LSFG.2 |
| 130 | LrMFG.4 | 129 | LrMFG.3 | 125 | LparsOp1 | 135 | LSFG.3 |
| 132 | LrMFG.6 | 133 | LSFG.1 | 126 | LparsOp2 | 136 | LSFG.4 |
| 138 | LSFG.6 | 141 | LSFG.9 | 131 | LrMFG.5 | 139 | LSFG.7 |
| 140 | LSFG.8 | 142 | LcMFG.1 | 135 | LSFG.3 | 142 | LcMFG.1 |
| 158 | LPostCinG.2 | 157 | LPostCinG.1 | 137 | LSFG.5 | 157 | LPostCinG.1 |
| 159 | LIstCinG.1 | 165 | LPostCG.6 | 144 | LcMFG.3 | 177 | LSPG.6 |
| 166 | LPostCG.7 | 180 | LIPG.2 | 152 | LPreCG.8 | 182 | LIPG.4 |
| 174 | LSPG.3 | 183 | LIPG.5 | 154 | LParaCG.2 | 188 | LPC.5 |
| 179 | LIPG.1 | 186 | LPC.3 | 170 | LSMG.4 | 201 | LFusG.2 |
| 203 | LFusG.4 | 201 | LFusG.2 | 173 | LSPG.2 | 217 | LSTG.1 |
| 209 | LITG.3 | 205 | LEntRG.1 | 194 | LLatOG.4 | 218 | LSTG.2 |
| 210 | LITG.4 | 212 | LMTG.2 | 200 | LFusG.1 | 220 | LSTG.4 |
| 211 | LMTG.1 | 217 | LSTG.1 | 204 | LPHG.1 | 221 | LSTG.5 |
| 213 | LMTG.3 | 219 | LSTG.3 | 208 | LITG.2 | 222 | LTransTG.1 |
| 223 | LIns.1 | 222 | LTransTG.1 | 220 | LSTG.4 | 231 | LAccu |
| 226 | LIns.4 | 225 | LIns.3 | 224 | LIns.2 | 233 | LAmy |

### Network-level differences in hubness and segregation of HFA-memory regions and neighboring control nodes (region-effect), and between patients and controls (group-effect)

At a network level, MEM had a lower AvgBC compared to CNs (MEM vs. CN.1--, F’s>14.16, FDR-p’s <3.4×10^-3^; MEM vs. CN.2 -- F’s> 20.16, FDR-p’s <2.8×10^-4^; MEM vs. CN.RAND -- F’s> 23.63, FDR-p’s <6.6×10^-6^. The df’s for each comparison=72). Neither the main effect of the group nor the interaction between group and region was significant.

Patients had a lower AvgCC compared to controls (Patient vs. Controls for MEM vs. CN.1: F’s> 8.2827, FDR-p’s<0.004; Patient vs. Controls for MEM vs. CN.2: F’s> 7.1628, FDR-p’s <0.0088; Patient vs. Controls for MEM vs. CN.RAND: F’s> 8.1457, FDR-p’s <0.007. The df’s for each comparison = 72). Neither the main effect of the region (MEM vs. CN.1, CN.2, and CN.RAND), nor the interaction between group and region was significant (Figure 3).

**Figure 3:**
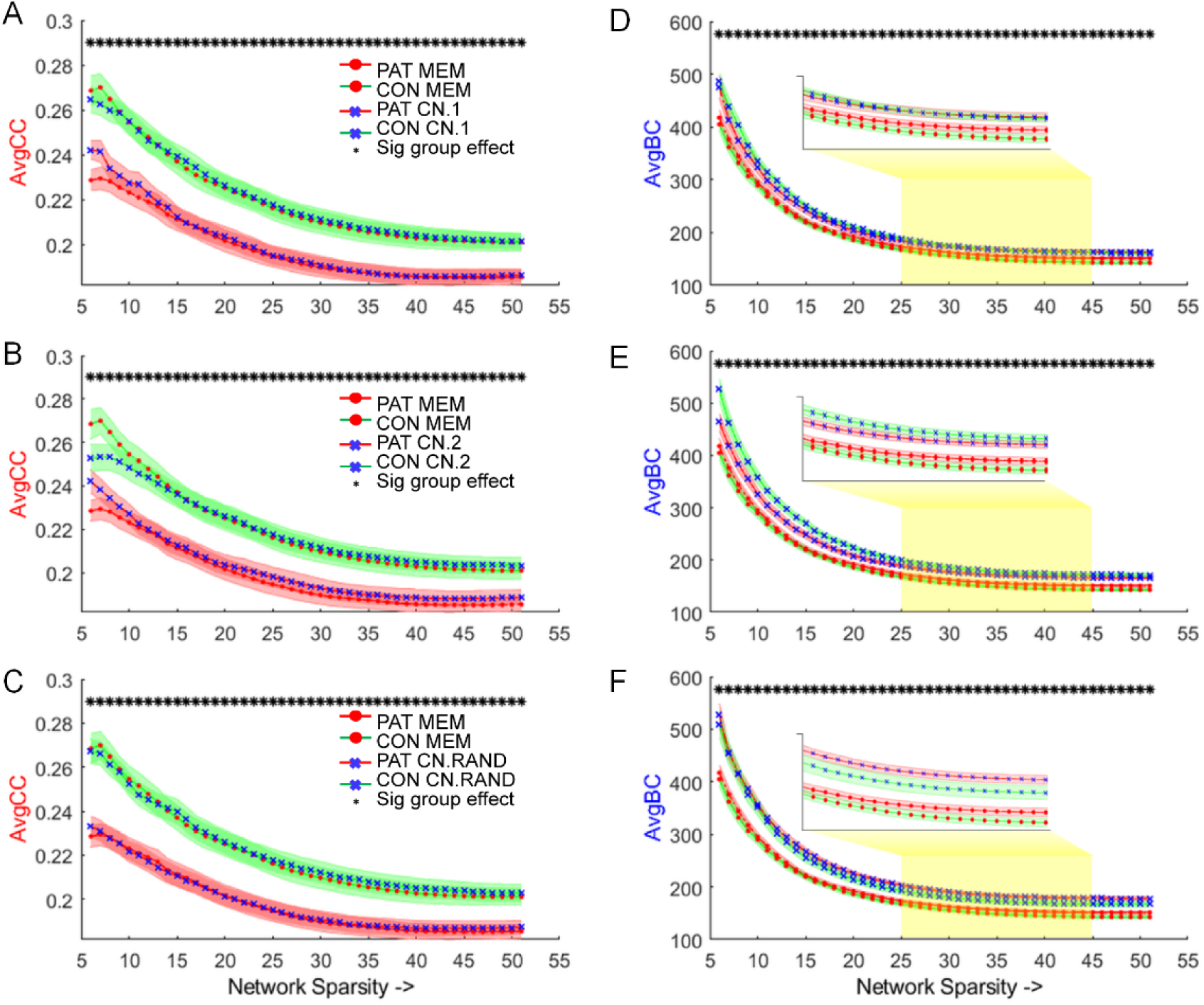
Network-level ANOVAs on AvgCC and AvgBC testing group-by-region effects. Composite network level (AvgCC and AvgBC) differences in MEM and CN (region-effect) between patients (PAT) and controls (CON) (group-effect). Panels A, B, and C show network segregation (AvgCC) was lower in patients compared to controls (* - indicates a significant main effect of the group). Panels D, E, and F show hubness (AvgBC) of the HFA-memory network (MEM) was lower compared to their neighboring control nodes (CN.1, CN.2, and CN.RAND) (* - indicates the significant main effect of the region). The * indicates that the main effects of AvgCC and AvgBC were significant across network thresholds (sparsity levels from 5-50%). The yellow inset provides a magnified view of the difference in the AvgBC in the mid sparsity range.

### Nodal-level differences in segregation and hubness of the HFA-memory regions and neighboring control nodes (region-effect), and between patients and controls (group-effect)

The above network-level analyses made clear that hubness differed between the MEM and CNs irrespective of group, while segregation differed between the groups irrespective of regional differences. Because such effects may hide differences at a nodal level. We applied similar ANOVAs to BC and CC estimated at the nodal level (NodalBC and NodalCC respectively). NodalBC was lower in MEM compared to CNs involving the left rostral middle frontal gyrus indicating a significant region effect. The main effect of group and the group-by-region interaction was not significant (Table 3).

NodalCC was reduced in patients compared to controls across regions involving left caudal middle frontal gyrus, left inferior parietal gyrus, left precuneus, right fusiform, and lateral occipital gyri of the MEM indicating a significant group effect (Table 3). Neither the main effect of the region nor the group-by-region interaction was significant.

**Table 3:** ANOVAs on NodalCC or NodalBC testing group-by-region effects. NodalCC showed significant group-effect (patients had a lower NodalCC compared to controls irrespective of whether they were MEM or CNs). NodalBC showed significant region-effect (HFA-memory regions had lower NodalBC compared to CNs). (Regions of interest (ROIs), Clustering coefficient (CC), Betweenness Centrality (BC), HFA-Memory regions (MEM), Controls Nodes (CN). (F statistic of ANOVA (Fs), df= degrees of freedom, p values of ANOVA corrected for false discovery rate (pFDR), R.lateraloccipital (RLatOG), R.fusiform (RFusG), L.inferiortemporal (LITG), L.superiorparietal (LSPG), L.rostralmiddlefrontal (LrMFG)).

| **MAIN EFFECTS** |  |  |
| --- | --- | --- |
| **NodalCC**<br>**(Significant group-effect: Patient vs. Control)** | **Fs (df)** | **FDR-p's** |
| RLatOG.3 | >5.9921 (72) | <0.0057 |
| RLatOG.4 | >4.7211 (72) | <0.0062 |
| RFusG.1 | >4.1904 (72) | <0.0277 |
| RFusG.2 | >4.5526 (72) | <0.0300 |
| LSPG.3 | >4.3929 (72) | <0.0044 |
| LITG.4 | >4.2426 (72) | <0.0136 |
| **NodalBC**<br>**(Significant region-effect: MEM vs. CN)** |  |  |
| LrMFG.4 | >4.4909 | 0.0004 |

### HFA-memory network decomposition among the established intrinsic networks

The HFA-memory network had a diverse “membership”, sharing only modest overlap with any single intrinsic network, involving a complex set of cognitive and sensorimotor processes involved in effective memory encoding. The HFA-memory network comprised 16% of TN, 12% of FPN, 14% of DAN, and 13% of COpN, amounting to a large 55% membership in cognitive control systems (Figure 4A). This stands in contrast to very limited membership of 15% with DMN, 6% with SMN, and 24% with VIN (Figure 4A). The HFA-memory network ranked 4^th^ (AvgBC: 200±42) against the 9 intrinsic networks and differed significantly from all but one in AvgBC (Figure 4B) (Appendix B, C).

**Figure 4:**
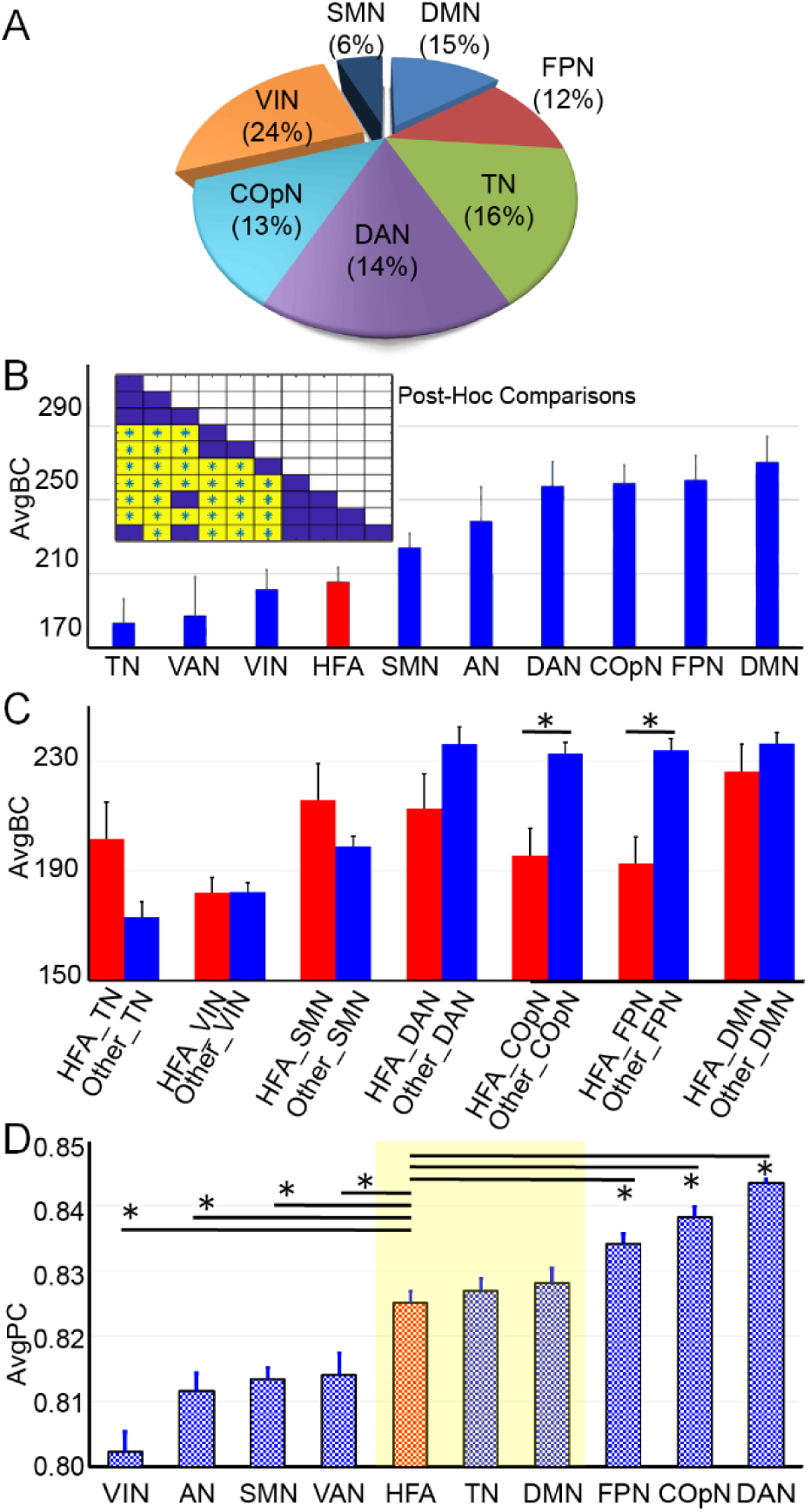
Nature of hubness of the HFA-memory network. Pie-graph shows percent contribution of each intrinsic network to the HFA-memory network. The HFA-memory network appears to be a diverse, composite network with contributions from other networks (B) HFA-memory network had a BC value distinct from all other intrinsic networks except the SMN (inset, post-hoc comparisons between the networks in the same order as the bar graph). DMN and the FPN, which form the core hubs in the brain had the highest AvgBC values. In fact, with the exception of the DMN, all the networks display BC values distinct from at least 5 of the other networks. (C) shows the difference in BC of HFA member ROIs of an intrinsic network compared to BC of the other nodes (non-HFA) in the same network [HFA_COpN vs. Other_COpN, t= −3.47, p= 5.2* 10-4 and HFA_FPN vs. Other_FPN, t= −3.63, 2.9* 10-4). (D) Shows the participation coefficient (PC) of the HFA-memory network in comparison with the different intrinsic networks. The highlighted yellow area shows the similarity of HFA-memory network to TN and DMN. Each black horizontal line indicates the networks that had a different AvgPC compared to HFA-memory network following a pairwise comparison. (* - FDR corrected P-value for alpha set at 0.05, AvgBC-Average betweenness centrality of each network, AvgPC-Average participation coefficient of each network, TN-temporal network, VIN-visual integrated network, SMN-sensorimotor network, DAN – dorsal attention network, COpN – cingulo-opercular network, FPN-frontoparietal network, DMN – default mode network, ‘HFA_’ prefix – nodes of an intrinsic network that are part of HFA-memory network, ‘Other_’ prefix – nodes of an intrinsic network that are not a part of HFA-memory network).

The BC of ROIs of an intrinsic network that contributed to the HFA-memory regions differed from those that did not form a part of the HFA memory network. We found reliable differences only for the FPN (HFA_FPN vs. Other_FPN: t=-3.63, df=1, p=2.9×10-4) and COpN (HFA_COpN vs. Other_COpN: t=-3.47, df=1, p=5.2×10-4), in each case revealing higher BC for the regions of intrinsic networks not part of the HFA-memory network (Figure 4C). Accordingly, the BC levels of HFA-memory regions involved in intrinsic networks (e.g., HFA_COpN and HFA_FPN) can be said to differ from the non-memory regions of these intrinsic networks (Other_COpN, Other_FPN). A two-way ANOVA on these significant difference in the two networks showed that while the main effect of regional HFA status was significant (HFA_FPN vs. Other_FPN: F=19.53, df=3, p=1.9 ×10-5; HFA_COpN vs. Others_COpN: F=12.56, df=3, p=0.001), neither the main effect of group (patient vs. control) nor the group-by-network interaction was significant (Figure 5). The major finding was that cognitive control networks have a higher BC compared to the HFA-memory and primary processing networks.

**Figure 5:**
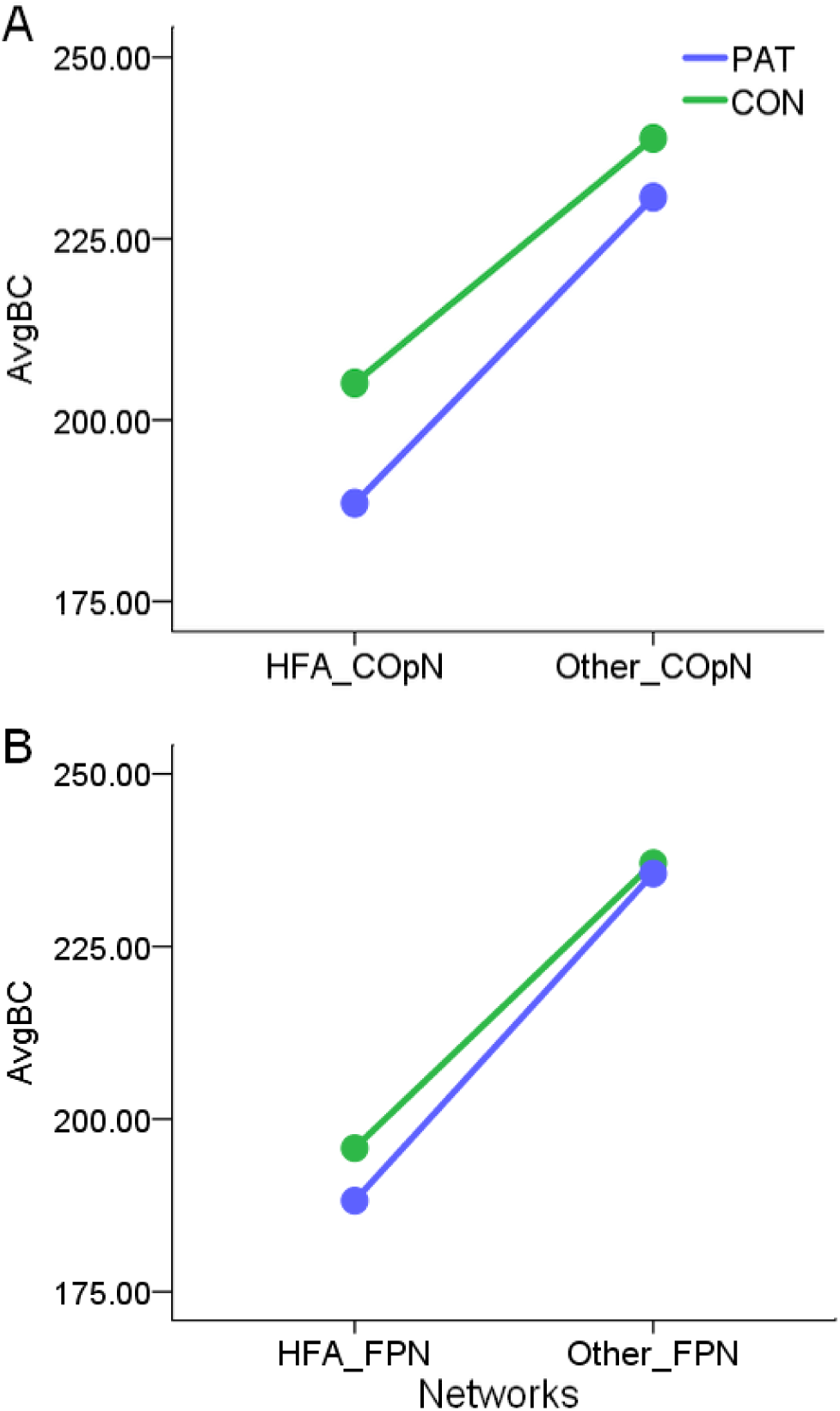
ANOVA comparing the effects of networks and subject groups for significant networks. The BC levels of HFA-members of intrinsic networks (HFA_COpN and HFA_FPN) differed from the remaining regions of a given intrinsic network (Other_COpN, Other_FPN). To determine if this effect was related to a patient vs. control group difference, we ran a two-way ANOVA. For both the CopN and FPN, the HFA-memory regions still differed from those regions not in the HFA memory network, even after accounting for the group difference. (A) The main effect of BC of HFA_COpN was lower than Other_COpN (F=12.56, p=0.001), while the main effect of group (patients vs. controls) was not significant (F=1.33, p=0.25), nor was the interaction between the group and HFA memory status (F=0.15, p=0.69). (B) The main effect of BC of HFA_FPN was lower than Other_FPN (F=19.53, p=1.9* 10-5), while the main effect of group (patients vs. controls) was not significant (F=0.27, p=0.65), nor was the interaction between the group and HFA memory status (F=0.09, p=0.76).

Inter-network interaction using PC revealed that the cognitive control networks and the HFA-memory network have higher AvgPC compared to the primary processing networks (Figure 4D, Appendix D). Importantly, the HFA-memory network possessed AvgPC levels similar to TN and DMN. The AvgPC of the cognitive control networks (FPN, COpN, DAN) was significantly higher than the HFA-memory network (Figure 4D, Appendix D).

### Multivariate machine learning to test the association of verbal-memory performance in epilepsy patients

As a final step, we tested if either hubness or segregation were relevant to an individual’s clinical memory performance. Though the two memory scores were clinically related (CVLT-TL, total recall across five trials and P.REC, the average percent recall across all trials), they were not collinear (r=0.27, p=0.12) and hence dealt with in two different RF models. The NodalBC of 4 MEM variables significantly predicted CVLT-TL performance (Figure 6A: NodalBC of ROI 1 and 2 of RFusG, LITG, and LSFG). The prediction response of this non-linear regression RF model was significant, as we found that actual CVLT-TL scores were correlated with the predictions made with the four MEM NodalBC predictors (r=0.8, p=2.04×10^-8^) (Figure 6A, inset). Among the 4 predictors, a stepwise regression model built with NodalBC of RFusG2 and LSFG6 emerged as significant (F=10.599, df=2, p=<0.0001: RFusG2: beta= -0.53, t=-3.7, p=0.001, VIF=1.52; LSFG6: beta=0.47, t=3.27, p=0.003, VIF=1.02).

**Figure 6:**
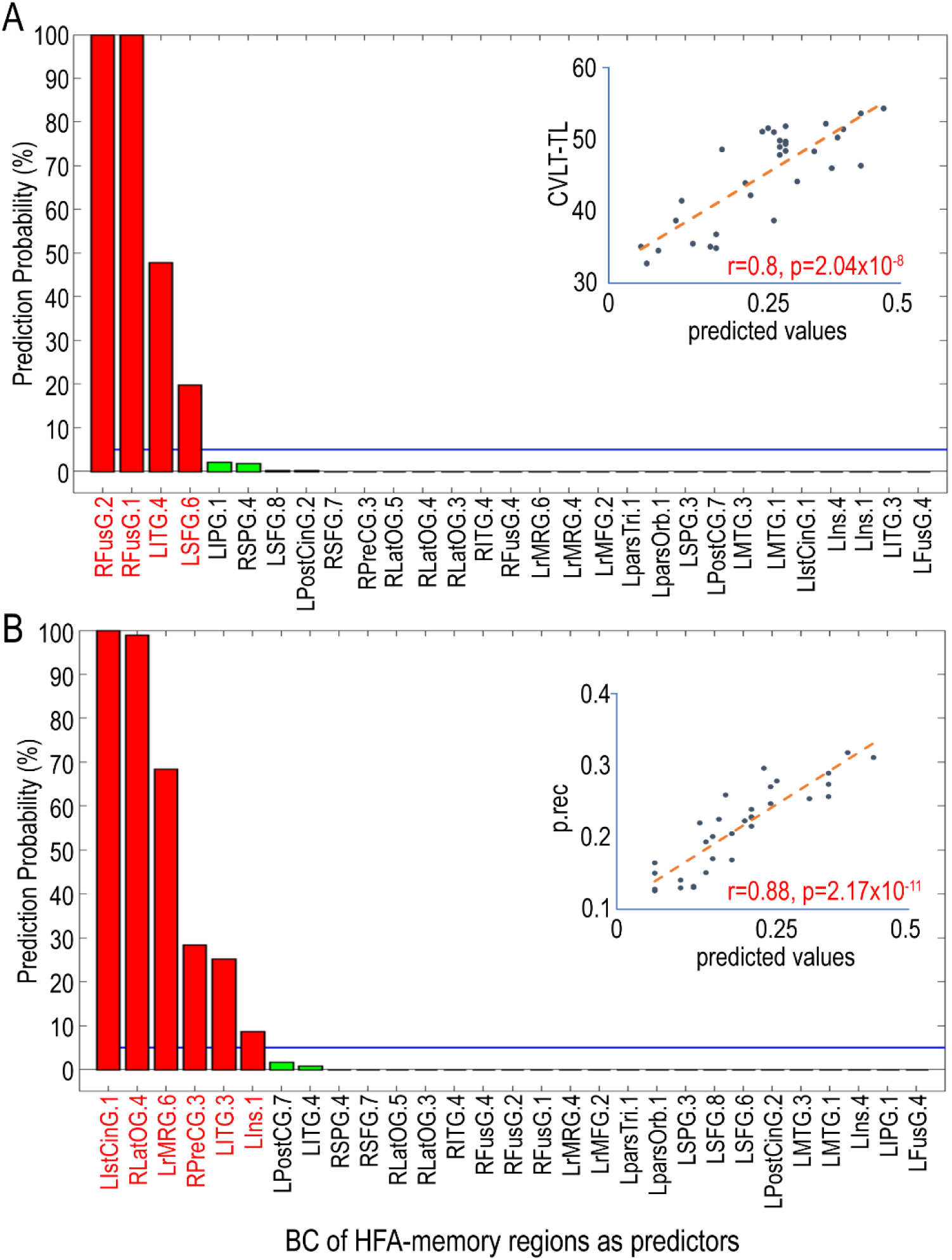
Random Forest Models to establish a relationship between hubness and verbal-memory performance. Random forest models help establish both linear and non-linear relationships between predictors and response. (A) BC of the HFA-memory regions (MEM) were used to predict CVLT-TL. The BC of the MEM were sorted in the descending order of their Variable Importance. LITG, LSFG, and RFusG were significantly associated with the CVLT-TL performance (inset-prediction of CVLT-TL performance made with BC of the MEM ROIs. (B) BC of the HFA-memory regions (MEM) was used to predict P.REC. The BC of the MEM were sorted on the basis of the probability of prediction based on 1000 permutations of the RF model. LIstCingG, LrMFG, LITG, LIns, RLatOG, and RPreCG were significantly associated with P.REC memory performance. Inset shows the prediction of P.REC performance during IEEG monitoring made with BC of the MEM ROIs. Measures exceeding beyond the blue line (the blue line indicates the p<0.05 threshold, below which none are considered significant). California verbal learning test (CVLT), number of times selected in the RF model (N), Betweenness Centrality (BC), High-Frequency Activity (HFA), correlation coefficient (r), probability value (p).

Similarly, the NodalBC of 6 MEM variables significantly predicted the P.REC performance (NodalBC of LIstCinG, LrMFG, LITG, LIns, RLatOG, and RPreCG) (Figure 6B). P.REC was correlated with the predictions made with these six MEM NodalBC measures (r=0.88, p=2.14×10^-11^; Figure 6B, inset), indicating that the RF model was accurate. A regression model testing the six predictors with P.REC produced no significant predictors.

In the RF models with the MEM NodalCC as predictors, none of the NodalCC measures were deemed significant in >95% of the permutations. Thus, none of these variables demonstrated a reliable ability to predict either P.REC or CVLT-TL performance. This modeling work established that resting-state hubness of the HFA-memory regions not only predicted the verbal-memory performance during the IEEG testing but also baseline levels of neuropsychological memory. Regional segregation showed no relation to either of these memory performances.

## DISCUSSION

A fundamental question in network neuroscience regards the specific relationship between intrinsic network architecture and task-defined regional brain activity. Thus, the goal of this study was to focus on the understanding of the broader network features that underlie and support memory functionality, particularly when such functionality is defined through a regionally limited and sparse technology such as IEEG. We demonstrated that by integrating spectral markers of memory encoding (HFA) with rsfMRI data, one can identify network features of an IEEG-derived verbal-memory network. This method also allowed us to reveal the diversity of memory encoding operations by revealing the contributions of multiple functional networks to successful memory. Given the inherent presence of a diseased sample when using IEEG data, we tested the generality of our findings by determining if the observed memory encoding network characteristics, as embedded in resting-state data, were similar in patients and healthy controls. To assess the clinical validity of these memory network characteristics, we tested whether these characteristics were associated with actual, baseline verbal-memory performance. In summary, with a combined IEEG-rsfMRI approach, we were able to lay out the network characteristics and multifaceted intrinsic cognitive components associated with successful verbal-memory encoding.

The data demonstrate that BC, a measure of hubness, as opposed to CC, a measure of segregation, is important for driving memory-encoding networks toward successful recall. Studies have indicated that in the human brain, the highest hub scores localize in the DMN and FPN (Power JD, et al., 2013;van den Heuvel MP et al., 2013). (Cole MW et al., 2013;van den Heuvel MP, et al., 2013), a finding consistent with our data. Compared to these dominant networks in the brain, we found that the HFA-memory network had a lower hub score, but nonetheless a level of hubness that sustained and supported successful memory encoding during IEEG testing. The apparent lower hub score for the HFA-memory network (i.e., AvgBC of MEM vs. CN), appeared to emerge from the fact that the control nodes with which they were compared were located in regions with hierarchically higher hub scores. While there have been studies showing the hippocampus to be a part of the brain’s hub architecture (Van Den Heuvel MP et al., 2011), there has been no study outside of this current data to show, one, the distinctiveness of the hub score (AvgBC) of a memory network and, two, it’s standing relative to the hubness values of other cognitive control networks. The hubness value of the HFA-memory network was not related to the presence of epilepsy and was obtained in the setting of normative baseline memory scores. As a point of contrast, we show that the patients and healthy controls differed with respect to the regional segregation (CC) of both the HFA-memory network and their neighboring control nodes. This segregation effect appeared to be a consequence of epileptic pathophysiology, a finding consistent with the previous literature (Haneef Z et al., 2014).

Nodal hubness and segregation mirrored the above composite network findings. During the estimation of memory encoding during IEEG testing, we excluded electrodes and regions that were a part of the clinically hypothesized epileptogenic zone in order to avoid the effect of epileptic interictal and ictal discharges on the estimation of HFA-memory regions, both in terms of affecting a patient’s ability to encode words, as well as the power changes related to the epileptic discharges in IEEG data. This is in accord with previous studies which have shown that significantly altered BC localized and lateralized to the epileptogenic regions in both IEEG and rsFC (Haneef Z, et al., 2014;Wilke C et al., 2011). Excluding those regions from analysis helped establish that the region-effects we observed at both the network level (AvgBC) and nodal level (NodalBC) were, indeed, associated with successful encoding. Note, in the setting of intact CVLT–TL performance, the appearance of regional AvgBC effects points to the likelihood that the various HFA-memory regions work in unison to maintain the integrity of verbal-memory encoding. The positive relationship we observed in the RF model between task-positive, cognitive control regions, and stronger memory performance is evidence of the way cognitive control benefits memory. The negative relationship seen between some NodalBC values and memory performance is less clear. On this point, there is literature showing that the areas involved in the early stages of memory encoding such as the ventral attention stream (i.e., RFUSG2) may be de-activated later in the memory encoding process (Burke JF, et al., 2015). Unfortunately, the low temporal resolution of rsfMRI would blur and be insensitive to this shift in activity, potentially leading to a negative correlation.

To clarify further the nature of the HFA-memory hubness findings, we compared the BC of the HFA-memory network to well-known intrinsic networks. In work utilizing network controllability measures, Gu and colleagues (Kauffmann L, et al., 2015) argued that the controllability of each of the intrinsic networks is unique. Applying this concept to the hubness results, we did see that the different cognitive networks operated at different hubness levels even in the resting-state among the same networks examined by Gu et al. 2015.

Importantly, we noted that the membership of different intrinsic networks in the HFA-memory network makes clear that the HFA-memory network utilizes a diverse set of intrinsic functionalities to drive successful verbal-memory encoding. The HFA-memory network has major membership (55%) from cognitive control systems that call upon attentional resources (DAN), lexical/semantic processing and memory consolidation (TN), and top-down control of executive functions (FPN, trial-specific control and selective attention; COpN, control of overall task goals and error monitoring) (Vaden KI, Jr. et al., 2013). Interestingly, we found that the HFA-memory regions that share membership with the intrinsic networks tend to operate at the same level of hubness as other constituents of the intrinsic network, with the exception of FPN and COpN networks (Figure 4c). These members of the HFA-memory network had a lower BC. (Sheffield JM et al., 2015). Other HFA-memory regions, utilizing cognitive processes that are part of other intrinsic networks, appear to operate at hubness levels that are comparable and optimal for the intrinsic networks as a whole (Figure 4c). Thus, our data indicates that effective memory encoding is not dominated by a central cognitive core, but is the result of a diverse set of componential functions distributed across multiple intrinsic networks, largely involving task-positive networks, all toward the goal of maximizing subsequent recall. This distribution across multiple regions is likely reflective of adaptive “associative” encoding, consistent with the extensive literature showing that recruitment of multiple cognitive and sensory processes is a crucial feature of an effective memory (Cowan N, 2017;Tulving E et al., 1996).

In order to go beyond a basic demonstration of the functional diversity of the HFA-memory network through data showing regional overlap, we investigated the interaction of the HFA-memory network with the different intrinsic brain networks. Participation coefficient analysis showed that regions of the HFA-memory network connect and interact with a diverse set of cortical regions and intrinsic networks, participating at a level higher than the primary processing networks (i.e., VIN, AN, SMN), comparable and equivalent to the other episodic memory networks (TN and DMN), but at a level lower than the cognitive control networks. The latter likely arises from the fact that cognitive control is applied to attention, language, visual-spatial, and motor functions, not just memory. Such data make clear that the HFA-memory network interacts intensively with both the other types of memory networks and the cognitive control networks, demonstrating that these interactions play a major role in achieving effective verbal memory encoding.

Lastly, we verified through RF modeling that the hubness of selected HFA-memory regions were significantly associated with both baseline verbal-memory performance (CVLT-TL) and successful recall during simultaneous IEEG-memory encoding (P.REC). The regions found to be predictive of memory performance are consistent with literature showing language and executive function processes (working memory, attention) mediate memory encoding, with such processes bringing semantic associations to bear during memory engram formation or utilizing a “central executive” to manipulate incoming information in working memory (Gutchess AH et al., 2005). Thus, similar to the data looking at the overlap with the intrinsic networks, the RF model pointed to the multivariate nature of memory encoding. Indeed, learning during the trials of the CVLT can be achieved through auditory attention and very short-term “holding” and covert verbal recitation of the information, i.e., good performance does not require long-term memory storage. The fact that this involved mostly left as opposed to right hemisphere regions could potentially point to a material-specific effect related to the word lists and covert verbal processing (Campoy G, 2008;Cowan N, 2008). The RF method produced a more sensitive and complex model of the relationship between hubness and memory performance, likely stemming from the fact that stepwise-regression only detects linear, not non-linear relationships. Overall, our data show that the hubness of regions matters more than segregation for the prediction of both baseline verbal-memory encoding abilities and successful recall following periods of HFA-associated memory encoding activity.

The two modalities we use, IEEG and rsfMRI, measure two biologically and technically unrelated signals (the direct activity of neuronal ensembles and metabolic-rate-driven synchronous fluctuations in cerebral-blood-flow, respectively), each based on very different spatio-temporal scales. There are multiple advantages of a method that uses IEEG HFA derived memory regions in individual subjects, and transforms them into a group-level whole-brain intrinsic functional connectivity map. First, from a statistical viewpoint, isolating HFA-memory regions allowed us to avoid, in both the inter- and intra-group comparisons, a large number of brain regions that are unassociated with memory encoding, reducing the chances of Type I error. The multiple comparisons performed in this study were restricted to the 29 regions with their corresponding controls regions, as opposed to correcting for ROIs of the entire Lausanne’s atlas. Second, the method of building a ‘Super-Brain’ allowed us to take advantage of the fine-grained temporal sequencing of cognitive events in IEEG, and reduce but not eliminate the problem of sparse spatial sampling. The use of a ‘Super Brain’ is widely established in cognitive studies to map reliable networks associated with memory encoding and retrieval (Greenberg JA et al., 2015;Kragel JE, et al., 2017). Third, IEEG is a spatially constrained investigative modality (i.e., sparse sampling). However, integration of the data with a modality such as rsfMRI allowed for evaluation of function throughout the brain even if patients were not implanted in those regions. This advantage has been exploited in previous studies involving isolation of the epileptogenic zone (Aghakhani Y et al., 2015), and in establishing task-related BOLD-gamma relations (Esposito F et al., 2013). Fourth, this type of integration opened up the door to studying the correspondence between BOLD signal changes and different frequency ranges, along with their linked behavioral activity. Indeed, linkages between neuronal synchrony and cognitive functions may be highly specific to the frequencies in which they occur. For instance, sensorimotor functions are regulated by beta synchrony (Brovelli A et al., 2004), and the correspondence between BOLD response and visual and auditory IEEG activity is regulated by alpha synchrony. Among the many different frequencies that can be studied in the IEEG, there has consistently been a close correspondence of the gamma band component of the local field potentials in the cerebral cortex with the BOLD signal for different motor, sensory and cognitive neuronal functions (Lachaux JP et al., 2007). Our results are certainly another example of this correspondence.

A handful of studies have performed network analysis directly on memory task-IEEG signals, studying the network synchrony present during encoding process (Burke JF, et al., 2013;Solomon EA, et al., 2017;Vecchio F et al., 2016). Such studies have emphasized different properties of the gamma frequency band-width, including spectral power changes, phase-locking value, and exact low-resolution topographical analysis (eLORETA). These studies have found that increased gamma spectral power, asynchronous gamma oscillations coupled with synchronous theta and increased gamma-smallworldness, were associated with better verbal and non-verbal episodic memory performance. Solomon et al., examined high gamma during verbal-memory and observed both synchronous and asynchronous activity. They found that regions in frontal, temporal, and medial temporal lobe cortex asynchronous with other regions, displayed a high level of connection strength, implying a high level of centrality or hubness within the memory network, a finding broadly consistent with the current results (Solomon EA, et al., 2017). We show that through repetitive processes of memory encoding, gamma frequency activity can establish synchrony to the point that even slow-moving resting-state BOLD fluctuations reflect their network impact. This network impact can be best characterized as establishing a level of hubness distinct from other intrinsic networks, yet spatially overlapping them, particularly those involved in cognitive control. In the first study of its kind, applying an integrated IEEG-fMRI method, we have shown that established markers of memory encoding correspond to meaningful network differences captured by rsfMRI. In clinical terms, knowing a region is a hub, playing an important role in multi-regional and multi-functional connectivity, may inform technologies trying to identify the most effective targets to electrically or pharmacologically stimulate for cognitive enhancement in areas such as memory (Ezzyat Y et al., 2017;Ezzyat Y et al., 2018;Kucewicz MT et al., 2018), or perhaps be used to identify the areas with sufficient influence over targets to generate effective neuro-feedback loops (Hohenfeld C et al., 2017;Murphy AC et al., 2017). Lastly, in the setting of clinical disorders such as epilepsy, knowledge of the broader functional network in which a given region is embedded, both its proximity to the epileptogenic (Horak PC et al., 2017;Towgood K et al., 2015), and the degree to which it operates as a core hub of the functional network, may contribute to the risk/benefit calculus of resecting a region, essentially allowing clinicians to better account for the network context and behavioral/cognitive impact of surgery.

In terms of the limitations of this study, the hippocampus was not significant for SME because it was found that in 11 of 37 subjects hippocampal electrodes were excluded from analysis as they were part of the epileptogenic zone, though they constitute an important part of the episodic memory network. Electrodes with higher epileptiform activity from regions such as the hippocampus, which is known to be involved episodic memory, were not included in the analyses, and the few electrodes that remained in these regions were not statistically significant for HFA. (Supplementary figure 8)(Yarkoni T et al., 2011). Note, the clinical memory performance of this cohort of patients was within the normal range making the point that while the hippocampus is a crucial structure in memory encoding, it’s actually the entire network associated with encoding that maintains the integrity of the function. We also want to acknowledge that there are a large number of network centrality measures, each with their own sensitivities to aspects of network architecture. Betweenness centrality has the assumption that short paths lengths are an important part of centrality, and that a region inter-connecting or “between” the separate modules is important, leaving open the question as to whether the BC hub itself is actually densely connected. Lastly, our capture of memory effects was limited to a specific band (44-100Hz). Given our data showing that memory-related asynchronous high frequency activity can be reliably mapped to resting state functional connectivity, future studies should focus on the whole brain network changes associated with other frequency bands, such as synchronous theta networks or high frequency activity greater than 100Hz and compare multiple bandwidths (i.e., not just high frequency but also, for example, low frequency theta synchrony) for their association with human memory (Khursheed F, et al., 2011;Solomon EA, et al., 2017). In some sense, our data can be seen as issuing an imperative regarding the importance not just of investigating and comparing multiple bandwidths for their association with cognition, but also to then integrate such findings with whole brain connectivity and networks so as to elucidate the full brain and cognitive impact of IEEG-defined regional functionalities.

In conclusion, the present study establishes that IEEG integrated with rsfMRI is a useful method for establishing the network characteristics associated with cognitive processes such as verbal-memory encoding, showing that repetitive gamma frequency activity can establish synchrony to the point that slow-moving resting-state BOLD fluctuations reflect their network impact. The HFA-memory network operates at distinct hubness levels and interacts with other intrinsic networks, all toward the goal of maximizing subsequent verbal recall. Importantly, our results clarify the conditions of the baseline, tonic state that supports effective, phasic, task-dependent memory encoding. This trait-like network feature holds true for both epilepsy patients and healthy controls. This diversity is potentially available as a source of compensatory reorganization in the setting of specific diseases such as epilepsy.

## Abbreviations

HFA: High-frequency activity
MEM: Brain regions showing HFA associated with verbal-memory encoding
CN: Controls Nodes
P.REC: Percentage of words recalled during IEEG memory testing
CVLT: California Verbal Learning Test
IEEG: Invasive Electroencephalography
BC: Betweenness Centrality
CC: Clustering Coefficient
PC: Participation Coefficient

## Author Contributions

Conceptualization and design of the study were by GC, JIT, and XH. The clinical evaluation and enrollment of patients in the EMU were by MS. The surgical procedures were headed and performed by AS. The neuropsychology data was collected and analyzed by NS, XH, and GC. IEEG and neuroimaging data were analyzed by GC, JK, WH, YE, and XH. Interpretation of results, drafting the manuscript and critical revisions were done by GC, XH, JK, WH, AS, MS, and JIT. JIT in the capacity of the corresponding author agrees to be accountable for all aspects of the work, ensuring the accuracy or integrity of any part of the work investigated and resolved.

## Acknowledgment

The authors thank the Epilepsy Monitoring Unit their help in data acquisition. We thank Dr. Dorian Pustina for his help with the machine learning strategy. The authors thank all the healthy controls and patients with epilepsy, whose data was used for this study. This work was supported, in part, by the DARPA Restoring Active Memory (RAM) program (Cooperative Agreement N66001-14-2-4032) to JK, YE, MS, AD, and JIT. JIT also acknowledges partial support from NIH 1R01NS112816-01. The views, opinions, and/or findings contained in this material are those of the authors and should not be interpreted as representing the official views or policies of the Department of Defense or the U.S. Government.

## References

Aghakhani Y, Beers CA, Pittman DJ, Gaxiola-Valdez I, Goodyear BG, Federico P (2015), Co-localization between the BOLD response and epileptiform discharges recorded by simultaneous intracranial EEG-fMRI at 3 T. Neuroimage Clin 7:755-763.

Alexander-Bloch AF, Gogtay N, Meunier D, Birn R, Clasen L, Lalonde F, Lenroot R, Giedd J, et al. (2010), Disrupted modularity and local connectivity of brain functional networks in childhood-onset schizophrenia. Frontiers in systems neuroscience 4:147.

Ashburner J, Friston KJ (2005), Unified segmentation. Neuroimage 26:839-851.

Axmacher N, Schmitz DP, Wagner T, Elger CE, Fell J (2008), Interactions between medial temporal lobe, prefrontal cortex, and inferior temporal regions during visual working memory: a combined intracranial EEG and functional magnetic resonance imaging study. J Neurosci 28:7304-7312.

Breiman L (2001), Random forests. Machine learning 45:5-32.

Brovelli A, Ding M, Ledberg A, Chen Y, Nakamura R, Bressler SL (2004), Beta oscillations in a large-scale sensorimotor cortical network: directional influences revealed by Granger causality. Proc Natl Acad Sci U S A 101:9849-9854.

Burke JF, Ramayya AG, Kahana MJ (2015), Human intracranial high-frequency activity during memory processing: neural oscillations or stochastic volatility? Curr Opin Neurobiol 31:104-110.

Burke JF, Zaghloul KA, Jacobs J, Williams RB, Sperling MR, Sharan AD, Kahana MJ (2013), Synchronous and asynchronous theta and gamma activity during episodic memory formation. Journal of Neuroscience 33:292-304.

Campoy G (2008), The effect of word length in short-term memory: Is rehearsal necessary? Q J Exp Psychol (Hove) 61:724-734.

Cole MW, Reynolds JR, Power JD, Repovs G, Anticevic A, Braver TS (2013), Multi-task connectivity reveals flexible hubs for adaptive task control. Nat Neurosci 16:1348-1355.

Cowan N (2008), What are the differences between long-term, short-term, and working memory? Prog Brain Res 169:323-338.

Cowan N (2017), The many faces of working memory and short-term storage. Psychon Bull Rev 24:1158-1170.

Esposito F, Singer N, Podlipsky I, Fried I, Hendler T, Goebel R (2013), Cortex-based inter-subject analysis of iEEG and fMRI data sets: application to sustained task-related BOLD and gamma responses. Neuroimage 66:457-468.

Ezzyat Y, Kragel JE, Burke JF, Levy DF, Lyalenko A, Wanda P, O’Sullivan L, Hurley KB, et al. (2017), Direct Brain Stimulation Modulates Encoding States and Memory Performance in Humans. Curr Biol 27:1251-1258.

Ezzyat Y, Wanda PA, Levy DF, Kadel A, Aka A, Pedisich I, Sperling MR, Sharan AD, et al. (2018), Closed-loop stimulation of temporal cortex rescues functional networks and improves memory. Nat Commun 9:365.

Friston KJ, Williams S, Howard R, Frackowiak RS, Turner R (1996), Movement-related effects in fMRI time-series. Magnetic resonance in medicine 35:346-355.

Geller AS, Schlefer IK, Sederberg PB, Jacobs J, Kahana MJ (2007), PyEPL: a cross-platform experiment-programming library. Behav Res Methods 39:950-958.

Genovese CR, Lazar NA, Nichols T (2002), Thresholding of statistical maps in functional neuroimaging using the false discovery rate. Neuroimage 15:870-878.

Greenberg JA, Burke JF, Haque R, Kahana MJ, Zaghloul KA (2015), Decreases in theta and increases in high frequency activity underlie associative memory encoding. Neuroimage 114:257-263.

Gutchess AH, Welsh RC, Hedden T, Bangert A, Minear M, Liu LL, Park DC (2005), Aging and the neural correlates of successful picture encoding: frontal activations compensate for decreased medial-temporal activity. J Cogn Neurosci 17:84-96.

Hagmann P, Cammoun L, Gigandet X, Meuli R, Honey CJ, Wedeen VJ, Sporns O (2008), Mapping the structural core of human cerebral cortex. PLoS biology 6:e159.

Haneef Z, Chiang S (2014), Clinical correlates of graph theory findings in temporal lobe epilepsy. Seizure 23:809-818.

He X, Bassett DS, Chaitanya G, Sperling MR, Kozlowski L, Tracy JI (2018), Disrupted dynamic network reconfiguration of the language system in temporal lobe epilepsy. Brain 141:1375-1389.

Hohenfeld C, Nellessen N, Dogan I, Kuhn H, Muller C, Papa F, Ketteler S, Goebel R, et al. (2017), Cognitive Improvement and Brain Changes after Real-Time Functional MRI Neurofeedback Training in Healthy Elderly and Prodromal Alzheimer’s Disease. Front Neurol 8:384.

Horak PC, Meisenhelter S, Song Y, Testorf ME, Kahana MJ, Viles WD, Bujarski KA, Connolly AC, et al. (2017), Interictal epileptiform discharges impair word recall in multiple brain areas. Epilepsia 58:373-380.

Humphries MD, Gurney K, Prescott TJ (2006), The brainstem reticular formation is a small-world, not scale-free, network. Proceedings Biological sciences / The Royal Society 273:503-511.

Jacques C, Witthoft N, Weiner KS, Foster BL, Rangarajan V, Hermes D, Miller KJ, Parvizi J, et al. (2016), Corresponding ECoG and fMRI category-selective signals in human ventral temporal cortex. Neuropsychologia 83:14-28.

Jin SH, Jeong W, Chung CK (2015), Mesial temporal lobe epilepsy with hippocampal sclerosis is a network disorder with altered cortical hubs. Epilepsia 56:772-779.

Kaiser M, Hilgetag CC (2006), Nonoptimal component placement, but short processing paths, due to long-distance projections in neural systems. PLoS computational biology 2:e95.

Kauffmann L, Bourgin J, Guyader N, Peyrin C (2015), The Neural Bases of the Semantic Interference of Spatial Frequency-based Information in Scenes. Journal of cognitive neuroscience 27:2394-2405.

Khursheed F, Tandon N, Tertel K, Pieters TA, Disano MA, Ellmore TM (2011), Frequency-specific electrocorticographic correlates of working memory delay period fMRI activity. Neuroimage 56:1773-1782.

Kragel JE, Ezzyat Y, Sperling MR, Gorniak R, Worrell GA, Berry BM, Inman C, Lin JJ, et al. (2017), Similar patterns of neural activity predict memory function during encoding and retrieval. Neuroimage 155:60-71.

Kucewicz MT, Berry BM, Kremen V, Brinkmann BH, Sperling MR, Jobst BC, Gross RE, Lega B, et al. (2017), Dissecting gamma frequency activity during human memory processing. Brain 140:1337-1350.

Kucewicz MT, Berry BM, Miller LR, Khadjevand F, Ezzyat Y, Stein JM, Kremen V, Brinkmann BH, et al. (2018), Evidence for verbal memory enhancement with electrical brain stimulation in the lateral temporal cortex. Brain 141:971-978.

Lachaux JP, Fonlupt P, Kahane P, Minotti L, Hoffmann D, Bertrand O, Baciu M (2007), Relationship between task-related gamma oscillations and BOLD signal: new insights from combined fMRI and intracranial EEG. Hum Brain Mapp 28:1368-1375.

Lega BC, Jacobs J, Kahana M (2012), Human hippocampal theta oscillations and the formation of episodic memories. Hippocampus 22:748-761.

Liaw A, Wiener M (2002), Classification and regression by random-Forest. Rnews 2:18-22.

Logothetis NK, Pauls J, Augath M, Trinath T, Oeltermann A (2001), Neurophysiological investigation of the basis of the fMRI signal. Nature 412:150-157.

Long NM, Burke JF, Kahana MJ (2014), Subsequent memory effect in intracranial and scalp EEG. Neuroimage 84:488-494.

Mizuhara H (2012), Cortical dynamics of human scalp EEG origins in a visually guided motor execution. Neuroimage 62:1884-1895.

Mizuhara H, Wang L-Q, Kobayashi K, Yamaguchi Y (2005), Long-range EEG phase synchronization during an arithmetic task indexes a coherent cortical network simultaneously measured by fMRI. Neuroimage 27:553-563.

Murphy AC, Bassett DS (2017), A Network Neuroscience of Neurofeedback for Clinical Translation. Curr Opin Biomed Eng 1:63-70.

Powell HW, Richardson MP, Symms MR, Boulby PA, Thompson PJ, Duncan JS, Koepp MJ (2007), Reorganization of verbal and nonverbal memory in temporal lobe epilepsy due to unilateral hippocampal sclerosis. Epilepsia 48:1512-1525.

Power JD, Schlaggar BL, Lessov-Schlaggar CN, Petersen SE (2013), Evidence for hubs in human functional brain networks. Neuron 79:798-813.

Rubinov M, Sporns O (2010), Complex network measures of brain connectivity: uses and interpretations. Neuroimage 52:1059-1069.

Rugg MD, Otten LJ, Henson RN (2002), The neural basis of episodic memory: evidence from functional neuroimaging. Philos Trans R Soc Lond B Biol Sci 357:1097-1110.

Sederberg PB, Gauthier LV, Terushkin V, Miller JF, Barnathan JA, Kahana MJ (2006), Oscillatory correlates of the primacy effect in episodic memory. Neuroimage 32:1422-1431.

Serruya MD, Sederberg PB, Kahana MJ (2014), Power shifts track serial position and modulate encoding in human episodic memory. Cereb Cortex 24:403-413.

Sheffield JM, Repovs G, Harms MP, Carter CS, Gold JM, MacDonald AW, 3rd, Daniel Ragland J, Silverstein SM, et al. (2015), Fronto-parietal and cingulo-opercular network integrity and cognition in health and schizophrenia. Neuropsychologia 73:82-93.

Solomon EA, Kragel JE, Sperling MR, Sharan A, Worrell G, Kucewicz M, Inman CS, Lega B, et al. (2017), Widespread theta synchrony and high-frequency desynchronization underlies enhanced cognition. Nat Commun 8:1704.

Solomon EA, Stein JM, Das S, Gorniak R, Sperling MR, Worrell G, Inman CS, Tan RJ, et al. (2019), Dynamic Theta Networks in the Human Medial Temporal Lobe Support Episodic Memory. Curr Biol 29:1100-1111 e1104.

Sperling MR, O’Connor MJ, Saykin AJ, Plummer C (1996), Temporal lobectomy for refractory epilepsy. JAMA 276:470-475.

Stam C, Tewarie P, Van Dellen E, Van Straaten E, Hillebrand A, Van Mieghem P (2014), The trees and the forest: characterization of complex brain networks with minimum spanning trees. International Journal of Psychophysiology 92:129-138.

Tewarie P, van Dellen E, Hillebrand A, Stam CJ (2015), The minimum spanning tree: an unbiased method for brain network analysis. Neuroimage 104:177-188.

Towgood K, Barker GJ, Caceres A, Crum WR, Elwes RD, Costafreda SG, Mehta MA, Morris RG, et al. (2015), Bringing memory fMRI to the clinic: comparison of seven memory fMRI protocols in temporal lobe epilepsy. Hum Brain Mapp 36:1595-1608.

Tracy J, Pustina D, Doucet G, Osipowicz K (2014), Seizure-induced neuroplasticity and cognitive network reorganization in epilepsy. Cognitive Plasticity in Neurologic Disorders:29.

Tracy JI, Lippincott C, Mahmood T, Waldron B, Kanauss K, Glosser D, Sperling MR (2007), Are depression and cognitive performance related in temporal lobe epilepsy? Epilepsia 48:2327-2335.

Tulving E, Markowitsch HJ, Craik FE, Habib R, Houle S (1996), Novelty and familiarity activations in PET studies of memory encoding and retrieval. Cereb Cortex 6:71-79.

Vaden KI, Jr., Kuchinsky SE, Cute SL, Ahlstrom JB, Dubno JR, Eckert MA (2013), The cingulo-opercular network provides word-recognition benefit. J Neurosci 33:18979-18986.

Van Den Heuvel MP, Sporns O (2011), Rich-club organization of the human connectome. Journal of Neuroscience 31:15775-15786.

van den Heuvel MP, Sporns O (2013), Network hubs in the human brain. Trends Cogn Sci 17:683-696.

van Diessen E, Diederen SJ, Braun KP, Jansen FE, Stam CJ (2013), Functional and structural brain networks in epilepsy: what have we learned? Epilepsia 54:1855-1865.

Vecchio F, Miraglia F, Quaranta D, Granata G, Romanello R, Marra C, Bramanti P, Rossini PM (2016), Cortical connectivity and memory performance in cognitive decline: A study via graph theory from EEG data. Neuroscience 316:143-150.

Wilke C, Worrell G, He B (2011), Graph analysis of epileptogenic networks in human partial epilepsy. Epilepsia 52:84-93.

Yan CG, Cheung B, Kelly C, Colcombe S, Craddock RC, Di Martino A, Li Q, Zuo XN, et al. (2013), A comprehensive assessment of regional variation in the impact of head micromovements on functional connectomics. Neuroimage 76:183-201.

Yan CG, Zang YF (2010), DPARSF: A MATLAB Toolbox for “Pipeline” Data Analysis of Resting-State fMRI. Frontiers in systems neuroscience 4:13.

Yarkoni T, Poldrack RA, Nichols TE, Van Essen DC, Wager TD (2011), Large-scale automated synthesis of human functional neuroimaging data. Nat Methods 8:665-670.

