## Supplementary material for "Tonic resting-state hubness supports high-frequency activity defined verbal-memory encoding network in epilepsy": SupplmentalData (from another version of this preprint: version 1, 2019-06-05)

### SUPPLEMENTARY DATA

### 1 Supplementary Methods

#### 1.1 Data Acquisition

The data collected during the study spanned both phase 1 and 2 of pre-surgical evaluation for epilepsy surgery and was aimed at localizing and lateralizing the epileptogenic lesion (EL), the irritative/interictal spiking zone (IZ) and the probable seizure onset zone (SOZ). Patients also underwent neuropsychological evaluation with the California Verbal Learning Test (CVLT), a test of episodic verbal learning and memory. The CVLT-Total learning measure (CVLT-TL) was utilized and it was calculated as the sum-total of all the words recalled from five separate learning trials (adjusted for age and years of education). This variable provided a measure of the patient's baseline verbal encoding ability.

Invasive EEG implantation (IEEG) and monitoring was carried out to precisely localize the IZ and the SOZ. The composite hypothesis regarding the epileptic focus derived from the phase-1 investigations help to determine the type (strips/grids/depth) and extent of electrode implantation to be undertaken (cortical surface/deep grey matter/mesial temporal and frontal structures, etc.). During the IEEG monitoring, all patients were administered a verbal episodic memory task (described below) synchronized with concurrent IEEG data recording.

#### 1.2 Resting-state functional MRI (rsfMRI)

All participants (patients and healthy controls) underwent rsfMRI scan and a T1 MPRAGE. For the rsfMRI scan, all participants were instructed to remain still throughout the entirety of the scan, keep their eyes closed, and not to move or fall asleep. Prior to starting the scan, the participants were familiarized to the MRI environment. The rsfMRI was acquired under strict environmental conditions regulating for head movement and noise reduction to ensure that the subtle effects of cognitive mapping being planned in the study were not affected by artifact. Low-noise MR acquisition was utilized to aid in the detection of subtle differences in neuronal networks (Rondinoni C et al., 2013). Significant noise reduction was obtained by providing noise cancellation ear buds and noise-reducing headphones. To prevent head-motion, foam pads were used to stabilize the head, while the headphones provided a restriction to any kind of head-motion within the coil.

#### 1.3 IEEG acquisition and free recall testing

Patients were implanted with IEEG electrodes [subdural, grids and strips (10mm resolution) or depth electrodes (5-10mm resolution)]. During IEEG monitoring, patients performed two variants of a delayed free-recall task (including either categorical or unrelated word lists; https://github.com/pennmem/wordpool) in three distinct phases of encoding, delay and retrieval (Kragel, Ezzyat et al. 2017). Participants were instructed to commit the list of words to memory and recall them in any order following a math-filled delay. The task was presented at the bedside using PyEPL software and the IEEG data was simultaneously recorded using Nihon-Kohden IEEG system [sampling rate >500Hz](Geller, Schlefer et al. 2007). Time-synced markers for stimulus presentation were recorded on the IEEG signal. During encoding, lists of 12 words were presented (each word for 1600ms followed by a blank inter-stimulus interval (ISI) of 750 to 1000ms). A 20s post-encoding delay period containing an arithmetic task was given to disrupt memory for end-of-list items (patients solved math problems of the form A+B+C=X using a response keypad). This was followed by the recall period of 30s, demarcated by a row of asterisks and an accompanying auditory tone (800 Hz, 300ms), during which the participant would recall as many words as possible from the most recent list presented, in any order. Recalled responses were digitally recorded and parsed/scored offline using the University of Pennsylvania Total Recall program (http://memory.psych.upenn.edu/TotalRecall). A measure of immediate recall, representing the percentage of words recalled across the multiple trials was utilized (percent recall, P.REC). Participants performed up to 25 recall trials (i.e., lists) in a single recall session. Supplementary figure 1 provides an illustration of the free-recall test session performed in the EMU.

##### Supplementary figure 1: Simultaneous IEEG acquisition and free recall testing

This is a schematic representation of simultaneous IEEG acquisition and free recall testing sessions administered to the epilepsy patients in the EMU. There were 3 phases of the task: 1) Encoding, where a list of new random words are presented to the subject and they are asked to commit them to memory, 2) Distractor task, involving a simple math problem to distract the subject from rehearsal or covert repetition of the words, and 3) Recall, where the subject is asked to recall the words presented during the encoding phase. This design provided information about IEEG activity during the words that were successfully memorized versus and those that were forgotten.

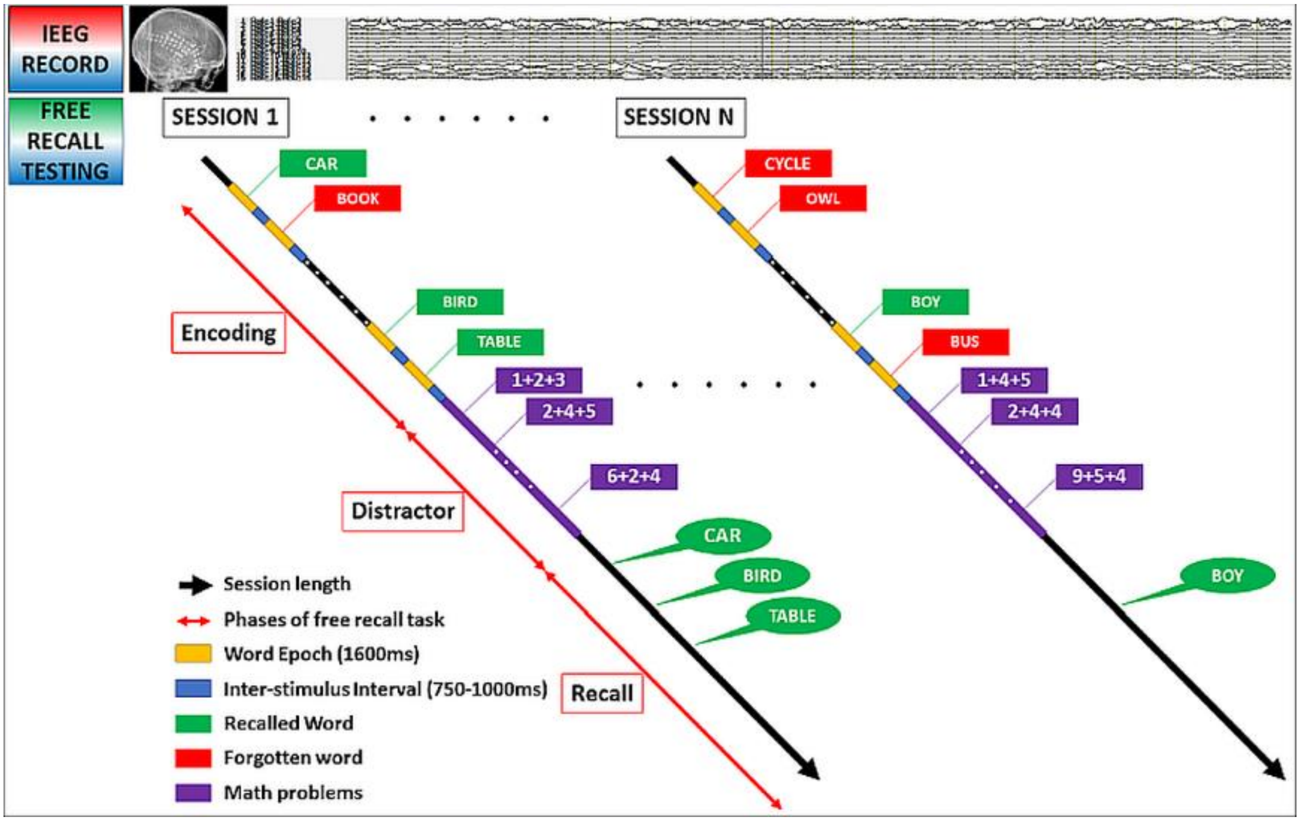

#### 1.4 Data Analysis

#### 1.5 Word encoding power analysis

The bipolar time-series derived from the implanted electrodes were bandpass filtered with a 4th order Butterworth band reject filter from 58 to 62Hz to exclude line-noise. To quantify memory-related changes in spectral power, complex-valued Morlet wavelets (wave number 6) were convolved with the down-sampled (200Hz) bipolar IEEG signals to obtain magnitude information. Ten wavelets with center frequencies logarithmically-spaced between 40 and 100Hz were used to estimate changes in HFA power. The average spectral power for each encoding event was computed from the onset of each stimulus for a duration of 1600ms, with a buffer of 1000ms. Prior to computing the subsequent memory effect (SME), power values for each bipolar pair were log transformed and normalized to have zero mean and unit standard deviation within each session. The SME was computed for each bipolar pair by computing an unpaired t-statistic on the distribution of power values associated with all successful and unsuccessful trials (n.b., unsuccessful trials were the words not recalled during retrieval).

#### 1.6 Functional MRI Preprocessing

All imaging data were preprocessed using Data Processing Assistant for Resting-State fMRI-Advanced Edition (http://www.rfmri.org/**DPARSFA**)(Yan CG and Zang YF, 2010), a MATLAB toolbox based on Statistical Parametric Mapping (http://www.fil.ion.ucl.ac.uk/spm/software/spm8). The order of preprocessing was as follows. The first three dynamics were discarded to maintain BOLD signal homogeneity. Following this, we performed a slice timing correction in an interleaved manner with the mid-time slice as a reference. Next, the T1-MPRAGE and the corresponding fMRI dynamics were re-oriented and realigned accurately to the anterior commissure-posterior commissure (AC-PC) plane in a subject-wise manner. Following brain extraction using Brain Extraction Toolkit (BET) (https://github.com/neurolabusc/MRIcro), the T1-MPRAGE images were co-registered to functional scans and then segmented along with neo-segmentation algorithm+DARTEL. A nuisance covariate regression was performed using Friston’s 24-parameter model (Friston KJ et al., 1996). The CSF and WM masks derived using segmentation were regressed out using CompCor (Ashburner J and Friston KJ, 2005;Yan CG et al., 2013). The resultant fMRI data were filtered between 0.008-0.1Hz. The images were normalized to the MNI space and smoothed using Gaussian Kernel of 4mm full width half maximum (FWHM). Bivariate ROI-to-ROI correlation matrices were extracted using Lausanne 234 ROI atlas for further graph theory analyses, testing for the association between the different brain parcels.

#### 1.7 Intrinsic Functional Connectivity mapping using Graph Theory Analysis

##### 1.7.1 Minimal Spanning Tree (MST)

MST of a graph is defined as the subgraph of an undirected graph that connects all nodes, with minimum total edge weight, without forming any loops. MST performs better on weighted undirected graphs (Tewarie P et al., 2015). MST is insensitive to the threshold and density value and is a good technique for graph binarization in the later analysis (Stam C et al., 2014). The absolute correlation of 234x234 ROI rsFC matrices was used to construct binary undirected graphs. MST-based networks contain the same number of connected nodes for all subjects, allowing for reliable group-level comparisons (van Diessen E et al., 2013). To do so, we first defined the MST that connected all 234 regional nodes with 233 edges for every subject (Alexander-Bloch A et al., 2013;AlexanderBloch AF et al., 2010). MST generates a highly-sparse network (MST network with N nodes has N-1 edges), and thus many short-range path connections (which is observed in many real systems) will be absent in MST. MST 'sparsely' represents a 'skeleton' of the brain network and it does not form clusters or loops at the regional level thereby precluding its use as a biologically meaningful sparse representation (Alexander-Bloch AF et al., 2010) . As a consequence, to obtain a series of sparse, yet fully connected biologically plausible graphs for each individual, additional edges were added to the MST in the descending order of the correlation values (Su L et al., 2013), yielding a series of networks with connection density ranging from 5% to 50% in increments of 1% (He X et al., 2017).

Two network topology measures were calculated using the Brain Connectivity Toolbox (www.brain-connectivity-toolbox.net)(Rubinov M and Sporns O, 2010): 1) clustering coefficient [CC] as a measure of brain segregation and 2) betweenness centrality [BC] as a measure of hubness. The CC of a node is the probability that all the neighbors of a node are also connected to each other, thereby representing the local segregation or "cliquishness" of the brain. BC refers to the relative importance of a brain region connecting two subnetworks. The BC of a particular node is the fraction of shortest geodesic paths in the network that pass through it. In the setting of this study, a difference in hub value can be interpreted as the degree of importance by which the nodes may vary between functionally relevant regions (HFA-memory versus surrounding control regions) or between our DRE patients and controls.

##### 1.7.2 Defining the HFA-memory regions (MEM) and control nodes (CN) for rsFMRI Analysis

Defining the HFA-memory regions (MEM) and their surrounding control nodes (CN), and integrating these ROI's with the rsfMRI data was the crux of the study. The properties of MEM at both the regional and network level were studied, utilizing regions that survived statistical thresholding for significant subsequent memory effects (SME). In this study, we focused on establishing network differences between MEM and CNs in rsfMRI. The rationale for using neighboring regions as control nodes was to avoid comparison to more distant regions of the brain, housing functions completely unrelated to the target HFA-memory regions.

##### 1.7.3 Assigning the bipolar electrode contacts to the different brain parcels

In order to assign the bipolar electrode contacts (derived as the mid-point of the two contacts) to different brain parcels, we proceeded with the following steps. Based on the electrode locations derived for each subject (Methods section of the manuscript, "anatomical localization of the bipolar electrode contacts"), we were able to identify the electrode locations in native coordinates (i.e., at individual subject level). These native coordinates were then normalized to MNI space by using the same methods described in previous studies by the co-authors (Burke JF et al., 2015;Kragel JE et al., 2017). All the MNI-normalized contact coordinates from the 37 subjects were pooled together for this procedure. The resultant 3731 IEEG bipolar contacts were superimposed onto one single brain, referred to as the "super-brain" (Figure 1). The electrodes were then assigned to each of the ROIs of the Lausanne atlas (234 brain parcels) using their 3D-Cartesian coordinates in Euclidian space (Hagmann P et al., 2008). In order to avoid the probability that the algorithm wrongly assigned the electrodes to subcortical structures, we excluded these regions, to have an atlas with only 222 implant relevant regions.

Each bipolar coordinate was warped to the coordinate of the nearest ROI of the Lausanne atlas. For this, we defined two matrices, vertices (V) and points (P). V is a Vx3 matrix of 3D Cartesian coordinates of a mesh generated from the Lausanne atlas and P is a Px3 matrix of 3D Cartesian coordinates of the 3731 IEEG contacts. Each of the coordinates in P was assigned to each of the ROIs by calculating their distance to the closest points in the vertex cloud V, within the volume size of each ROI of the atlas. This was calculated as the shortest Euclidean distance (ED) between the coordinates in V and P such that the vertex in P was a point in V. The function then warps the electrode coordinates to the nearest template coordinate of V (minimum/shortest ED) in the cloud and repeats this for all the coordinates in P. This allowed us to group the 3731 contacts into 222 ROIs. Once this was done, the t values of the multiple electrodes within each ROI, representing the strength of SME (the power difference in the high gamma frequency between words that were encoded vs those words that were not encoded) were submitted to a positive tailed one-sample t-test to determine if that ROI as a whole contained significant levels of memory-related HFA. In supplementary figure 2, we provide an example of the assignment of the implanted electrodes to a parcel (ROI) involving the primary motor cortex.

**Supplementary figure 2: Assigning the IEEG bipolar derivations to the different brain parcels**

This figure displays an example of the assignment of the coordinates of the IEEG bipolar derivations to the 3D coordinates of the nearest ROI of the Lausanne's atlas based on their Euclidean distance proximity. We notice that a total of 188 electrodes (green spheres) were placed in bilateral precentral gyri (yellow regions) in the 37 patients included in this study. Using this method of assigning electrodes, we were able to reliably assign the 3731 electrodes to the 222 ROIs of the modified Lausanne's atlas.

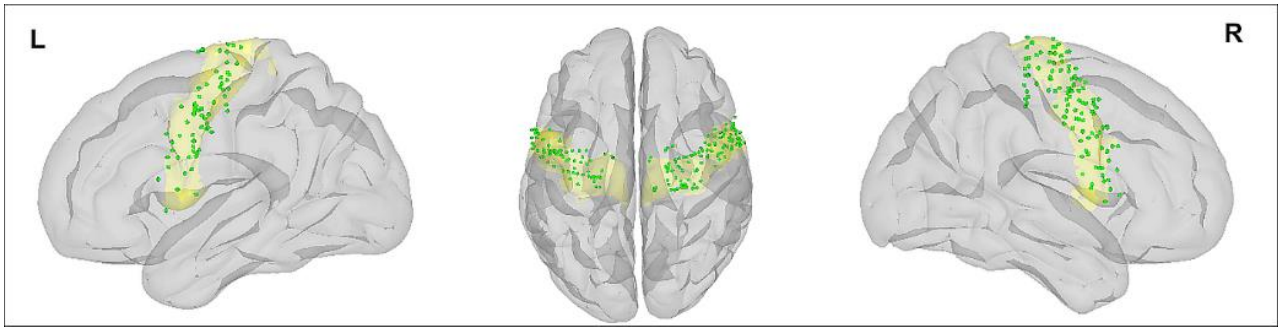

**Supplementary figure 3: The three control nodes (CN) in relation to the MEM ROIs**

Control Nodes (CN) involved parcels neighboring the HFA-memory regions (MEM). Three different types of CN were identified based on Euclidean distance proximity to the vertex of the nearest cortical ROI of the Lausanne 234 atlas. CN.1 (red) was the closest parcel to a given MEM ROI. CN.2 (blue) was the second closest, and, finally, a random ROI chosen among the 5 parcels closest to a given MEM ROI was designated as CN.RAND (cyan).

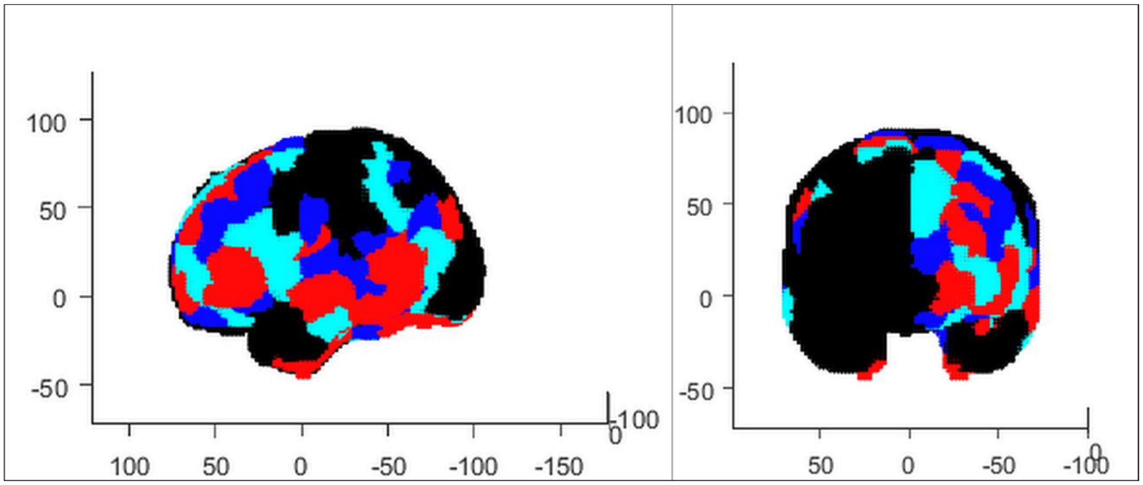

#### 1.8 ANOVA model for Average CC and BC, also Nodal CC and BC analyses:

A mixed model ANOVA on the graph theory measures was conducted at two different levels: 1) ‘composite network’ level (AvgCC and AvgBC) and 2) ‘individual region’ level (Nodal CC and Nodal BC). Each ANOVA contained one between-subjects factor (group-effect-Patients vs Controls) and one within-subjects factor (region-effect, repeated-measure with two levels, MEM and Control nodes). The AvgCC and AvgBC measures averaged values across the separate MEM (37 regions) and CN regions (37 values). The Nodal CC and Nodal BC measures were computed at the ROI level. This yielded 37 ANOVA’s involving the MEM ROI’s and the CN.1, CN.2, and CN.RAND, respectively, FDR corrected for each set of ANOVA’s. The group and region main effects and their interaction were included. Reliable effects required demonstration of significance across ANOVA’s involving all three CN’s after submission to

Boolean analysis. The final nodal ANOVA model utilized the average value across the relevant CN Nodal values.

##### 1.8.1 Boolean Conjunction Analysis:

In order to identify the significant MEM ROIs, we used Commutative Boolean Conjunction analysis. Only MEM ROIs that significantly differed in comparison to CN.1, CN.2, and CN.RAND across all three of these ANOVA models were considered and used in subsequent analyses. This is a stringent requirement for significance, indicating that a particular MEM ROI demonstrated a reliable graph property difference (BC or CC) compared to two proximal ROI's (CN.1, CN.2) and one random ROI (CN.RAND) (Equation 1, supplementary figure4).

*Sig region-effect = (Sig MEM from ANOVA.1) ʌ (Sig MEM from ANOVA.2) ʌ (Sig MEM from ANOVA.3)* - - - - - - - - - *Eq. 1*

Where Sig region-effect is the significant region-effect, ANOVA.1 is tested between MEM and CN.1, ANOVA.2 is tested between MEM and CN.2 and ANOVA.3 is tested between MEM and CN.RAND and ʌ is the commutative conjunction operator.

**Supplementary figure 4: Commutative Boolean Conjunction Analysis**

Comparison of the BC of the HFA-memory regions (MEM) with different control nodes (CNs) was done in order to test which of the regions showed a significant difference in their hub score. A different test if the Commutative Boolean Conjunction Analysis (Equation 1) graphically depicting a significant MEM Difference compared to all three control regions, CN.1, CN.2, and CN.RAND.

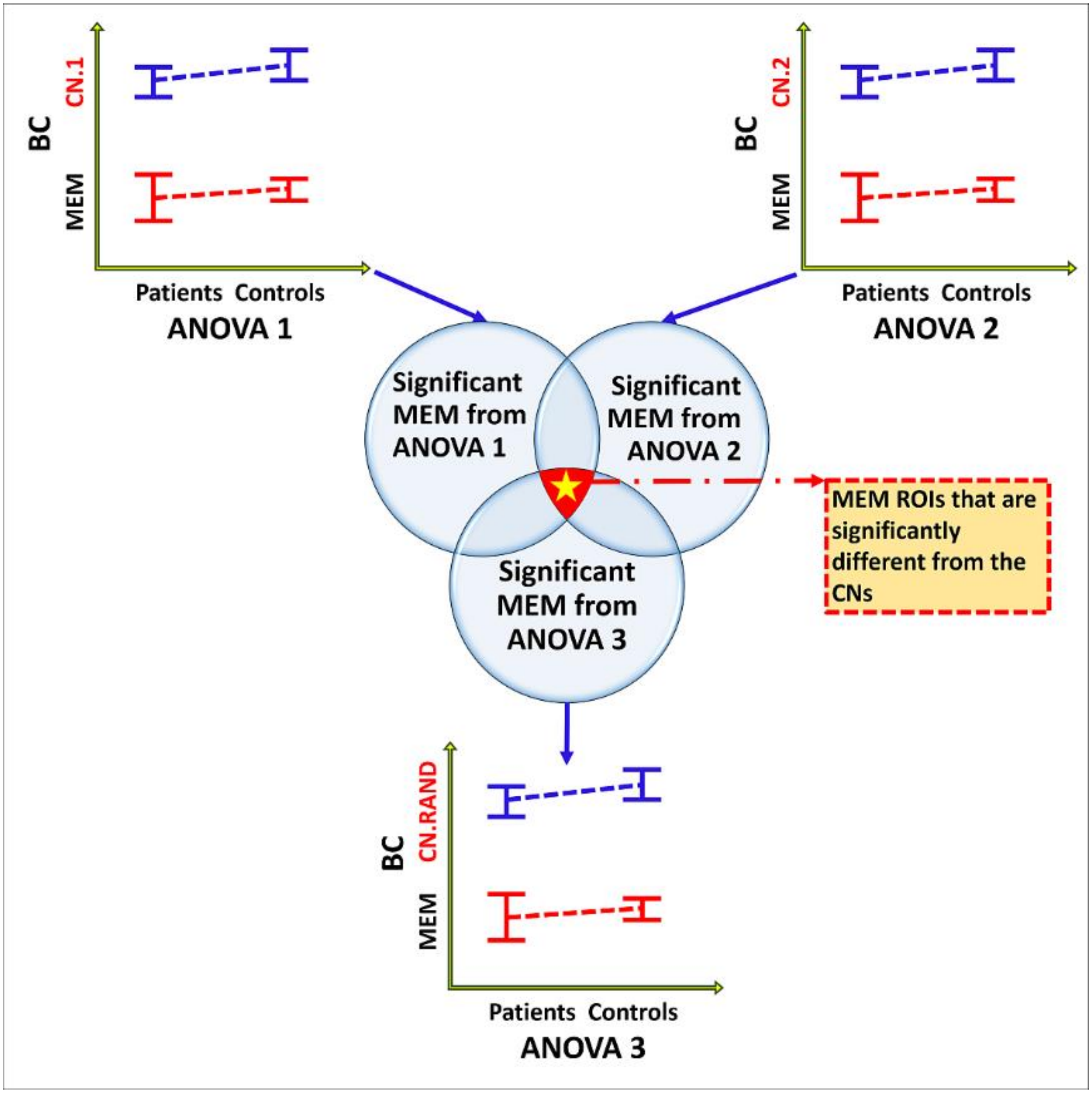

#### 1.9 Multivariate Machine Learning to Predict Free Recall Performance:

Using random forest we first ranked the fMRI graph indices (i.e., predictors) using VI, to identify the most relevant. Since multi-collinearity among predictors can bias VI (Tolosi L and Lengauer T, 2011) and alter predictor importance ranking, we applied a permutation-based method to statistically identify the most important features. Prior work has demonstrated that this approach is able to overcome bias from multi-collinearity (Altmann A et al., 2010). Briefly, we estimated the true VI for each predictor using random forest models with 1000 trees for 1000 repetitions to control for the variance introduced by bootstrapping. Then we established a null distribution of VI for every predictor by estimating their VI from the models trained with randomly permuted responses (i.e., CVLT-TL and P.REC in separate Random Forest models) for 1000 times. For each null distribution, a normal cumulative distribution was estimated based on its mean and standard deviation over the 1000 times. Accordingly, for the true VI of each predictor from each repetition, the probability (P statistics) can be estimated based on its corresponding normal cumulative distribution. We considered a VI as significant if its probability was larger than 95% (i.e., P = 0.05, one-tailed test), and the predictor was considered significantly relevant only when it was deemed significant 95% of the times (i.e., 950 out of 1000 permutations). Next, with the predictors found significant, we established RF models to predict the response. Because the bootstrap nature of the random forest approach can cause a slight variance in the predicted value across instances, we carried out 1000 repetitions for this procedure, and then used the average value as the final predicted response. Only a positive association between the predicted response and the true response would indicate that the latter could be explained by the information embedded in the predictors. Hence, we used a right-tailed Pearson correlation to test for an association.

While the RF model establishes the significant predictors, and the strength of the model prediction with actual performance, the direction of the association between the response and any given predictor is not given by the RF. To determine this we implemented a stepwise multiple regression model (SMR) on the response variable, using the significant RF model predictors as independent variables. This technique establishes the direction of the predictor relative to the response, and quantifies the strength of the association between the predictors, as a whole, and the response, as well as a relative ranking of the individual predictors, removing weak predictors from the model. More specifically, SMR picks the independent predictor with the highest ‘r’ value in the semi-partial correlation matrix and tests it in the regression equation (Equation 2).

$$Y = \beta_0 + \beta_1 x_1 + \beta_2 x_2 + \ldots\ldots \beta_n x_n + \varepsilon \text{ - - - - - - - - - - Eq. 2}$$

where, Y = response, $\beta_0$ = Y intercept, $x_1$ = first predictor, $\beta_1$ = weight of $x_1$, $\varepsilon$ is the random error. It subsequently picks the next predictor with the second highest 'r' value controlling for the variance from the first independent predictor. This process of adding more predictors to the equation is terminated when all the predictors have been included in the equation or no statistically significant improvement in the model 'R-squared' can be obtained with any of the remaining predictors. In this final model, the beta weights and their direction reflect the strength and direction of the relationship of each predictor with the response. One of the most important requirements of SMR is that no correlation be present among the independent predictors (known as collinearity). Accordingly, collinearity diagnostics are computed, as well as the variance inflation factor (VIF). Note, in our data both of these diagnostic factors were within normal ranges.

### 2 Results

#### 2.1 Participants:

A total of 37 patients were recruited for the study. Their complete clinical profile including chronological age, age of onset of epilepsy, gender, seizure types, MRI pathology lateralization and localization, interictal and ictal scalp EEG findings, site of IEEG implantation, IEEG localization of the electrical abnormality and the subsequent surgical procedure undertaken (if any) are detailed in **Supplementary Table 1**.

**Supplementary Table 1: Clinical profile of the DRE patients included in the study.**

| # | Age at study (Age at seizure onset), yr | Gender | AEDs at study | Seizure types | Anatomical MRI findings | Interictal Scalp EEG | Ictal scalp EEG | IEEG placement | IEEG findings | Surgery |
|---|---|---|---|---|---|---|---|---|---|---|
| 1 | 28.5(7) | F | LTG, LVA, DZP, CZP | FIAS, BTCS | R F DVA | Nil | R F onset | R F, L F | R F motor cortex | R F parasagittal resection |
| 2 | 34.5(32) | F | PGB, TPM | FAS, FIAS, BTCS | N | B/L FT spikes, sharp and slow waves | R midT onset | R FTP, L T | R subT onset | Extended right ATL |
| 3 | 24.7(20) | M | LVA | FAS, FIAS, BTCS | N | R T spikes | R T onset | R TF, L TF | R T neocortical onset | R ATL AH |

| | | | | | | | | | | |
|---|---|---|---|---|---|---|---|---|---|---|
| 4 | 49.3(40) | F | LTG, PHT | FAS, FIAS, BTCS | L F cavernous angioma | L FT sharp waves | L antT onset | L FTP, L Ins | L F and Ins onset | Cavernoma resection and insular focus resection |
| 5 | 20.6(16) | F | VPA, LCM | FIAS, BTCS | B/L PNH | B/L T spikes | Unclear | R F, L HG FT | L T widespread onset | Ablation of heterotopia |
| 6 | 28(19) | F | LVA, OXC, | FAS, BTCS | R F DVA | L FC spikes | Unclear | L F | L F onset | left frontal lobe resection with MST |
| 7 | 28.2(20) | M | LVA, LCM | FIAS | L med-post FP gliosis, R MTS | R T spikes, sharp waves | R neocortica lT onset | L postF, R T | R T onset | R ATL AH |
| 8 | 20(15) | F | VPA, LCM | FIAS, BTCS | N | B/L T sharp waves | R ant-midT onset | R FT, R TO | 2 foci: R subT onset and R orbF onset | R ATL AH + neocortical resection |
| 9 | 44.6(36) | F | LCM | FAS, FIAS, BTCS | L FT gliosis | L T slow waves | B/L independe nt T onset | L TF, R T | L T onset | L ATL+AH |
| 10 | 48.6(13) | M | TPM, LVA | FIAS, BTCS | N | B/L Tp sharp waves | Unclear R T onset | R FT, L FT | R F onset | right frontal and parietal resection |
| 11 | 44.9(40) | M | LVA | FAS, FIAS, BTCS | L Amyg bulky | L T spikes | B/L L > R T onset | B/L FT | B/L L > R independent tempral onset | RNS |
| 12 | 50(30) | M | LTG, CBZ, LCM | FIAS, BTCS | N | B/L T spikes and sharp waves | R antT onset | L FT, R FT | R T onset | R ATL AH |

| | | | | | | | | | | |
|---|---|---|---|---|---|---|---|---|---|---|
| 13 | 36.4(20) | M | CBZ, PHT | FIAS, BTCS | N | B/L T spikes | B/L - L > R T onset | B/L FT | B/L L > R independent tempral onset | No surgery |
| 14 | 44(3) | F | LTG | FAS, FIAS | L MTS | L T spikes | B/L - L > R T onset | B/L T | B/L subT onset | L ATL+AH |
| 15 | 52.1(33) | M | LVA, LCM | FIAS, BTCS | R > L MTS | Nil | L T onset | B/L T and Cin | B/L R > L independent tempral onset | R ATL AH |
| 16 | 38(10) | M | PGB, LTG | FIAS, | N | L TO spikes, sharp waves and polyspikes | L O theta rhythm | L FTPO and Ins (extensiv e) | L TO, L subT onset | No surgery |
| 17 | 18.7(13) | F | LVA, CZP, GBP | FAS, BTCS | R T hyperintensity | Nil | Central ryhthmic beta | R FP, R Ins, L FP | R SMA onset | resection R SMA |
| 18 | 39.2(17) | F | ZNS | FAS, FIAS | N | R midT spikes | Unclear R T onset | R HG Amy TO | R antT onset | R ATL AH |
| 19 | 27.8(12) | M | OXC, ZNS | FIAS, BTCS | N | Unclear F spikes | Unclear F onset | B/L F, R orbF | R F onset | Right frontal lobectomy |
| 20 | 33.9(20) | M | LVA, OXC | FAS, FIAS | N | Nil | Unclear L SMA onset | R > L FP, R T | R hemisphere motor and SMA onset | RNS |
| 22 | 49(47) | F | LVA, LTG,TP M | FAS, FIAS | N | L T spikes | L subT onset | L HG Amy entorhina l cortex | L ant-med T anset | Left ATL |
| 23 | 47.7(21) | F | ZNS, OXC | FIAS, BTCS | L F-Oper-Ins Encephalomal acia + L MTS | L T spikes | L TF onset | L FT Ins and Cin | L T onset | L ATL+AH+fron |

| | | | | | | | | | | |
|---|---|---|---|---|---|---|---|---|---|---|
| | | | | | | | | | | tal operculectomy |
| 24 | 40.6(32) | F | LCM | FIAS, BTCS | N | L TIRDA, L Tp sharp waves | L T onset | L FTP, R FT | L T onset, R T electrographic seizures | L ATL+AH |
| 25 | 39.2(6) | F | ZNS, LTG, PB, CZP | FIAS, BTCS | L T-Ins Gliosis with ex-vacuo dilatation of the ventricles | L T sharp | L T onset | L FT Ins | L Amy, HG, and L ant-inf Ins | Left ATL + Left F-OrbF corticectomy |
| 26 | 34.7(19) | F | TPM, OXC | FIAS, BTCS | N | L T sharp waves | L T onset | L FTO Ins | L HG sub T sub O | L ATL |
| 27 | 31.3(15) | M | VPA, PHT | FAS, BTCS | N | R ant-midT slowing and spikes | R FT onset | R FT subT TO | widespread R TF neocortical onset | No surgery |
| 28 | 25.2(12) | F | LTG | FIAS, BTCS | N | R T polymorphic delta, R antT sharp waves | L > R T onset | B/L FT | L T onset | L ATL |
| 30 | 42.7(27) | M | ZNS, OXC | FIAS | N | B/L T sharp waves | B/L independent L > R T onset | B/L T | B/L independent L > R T onset | No surgery |
| 31 | 56.2(50) | M | LVA, PGB, LZP | FAS, FIAS, BTCS | N | L T sharp waves | L T pole onset | L HG T orbF | L antT onset | LiTT |
| 32 | 48.5(35) | F | OXC, TPM | FIAS, BTCS | N | R T sharp wave, R FTC sharp waves, L | R ant-midT onset | B/L FT | R dorsolateral and F pole onset | R frontal lobectomy |

| | | | | | | | | | | |
|---|---|---|---|---|---|---|---|---|---|---|
| | | | | | | midT spikes | | | | |
| 33 | 23.2(4) | M | LVA | FAS, BTCS | PRES | Unclear | L T onset | L TP Ins Cin | L HG and post Amy | L ATL+AH |
| 34 | 21.1(0.1) | F | ZNS, LTG | FIAS | L HG Atrophy | L postT spikes | R F onset | B/L F T | No Seizures | No surgery |
| 35 | 25.6(19) | M | LVA TPM | FIAS | B/L Periventricular Nodular Heterotopia | L T spikes | L T onset | L TP, R T | L deep T heterotopia onset | GKRS - left-sided deep temporal heterotopia |
| 36 | 45(30) | F | LVA LTG | FIAS, BTCS | N | B/L independent T spikes | L neocortical T onset | B/L F T | B/L independent T onset | No surgery |
| 37 | 40.4(3) | F | Pr, CBZ, VPA | FIAS, BTCS | N | R mid-post T spikes | R mid-postT onset | B/L F T | L F onset | No surgery |
| 38 | 51.5(38) | F | LCM, ZNS | FIAS | L med-basal antT Encephalocoel | L FT spikes | L ant-midT onset | L HG Amy Ins Cin orbF postT TPO, R medT | L F onset | L ATL |
| 39 | 30.3(1) | M | LVA CBZ | FIAS, BTCS | R PO Heterotopia | R PO spikes | R PO onset | R FTPO (extensive) | R deep P onset | right parietal resection |

#-patient number; yr-years; F=Female; M-Male; PGB-Pregabalin; LVA-Levetiracetam; VPA-Valproate; LCM-Lacosamide; TPM-Topiramate; CBZ-Carbamazepine; ZNS-Zonisamide; OXC-Oxcarbazepine; Pr-Primidone; PHT-Phenytoin; CZP-Clonazepam; DZP-Diazepam; GBP-Gabapentin; PB-Phenobarbitone; LZP-Lorazepam; FAS-Focal Aware Seizures/Simple Partial Seizures; FIAS-Focal Impaired Awareness Seizures/Complex Partial Seizures; BTCS-Focal to bilateral tonic clonic seziures; N- Normal; R-Right; L-Left; B/L-Bilateral; F-Frontal; T-Temporal; P-Parietal; O-Ocipital, C-Central; Tp-Temporopolar; HG-Hippocampal gyrus; Amy-Amygdala; Oper-Opercular; Ins-Insular; Cin-Cingulate; Orb-orbital; SMA-Supplementary Motor Area; ant-Anterior; post-Posterior; mid-middle; med-medial; sub-subjacent; DVA- Developmental Venous Anomaly; PNH-Periventricular Nodular

Heterotopia; MTS-Mesial Temporal Sclerosis; PRES-Posterior Reversible Encephalopathy Syndrome; TIRDA-Temporal intermittent rhythmic delta activity;

#### 2.2 Electrode distribution:

In order to build the super brain, it was essential to determine the number of bipolar electrode contacts and their placement across the different lobes. Bipolar contacts were excluded from the analyses if there were artifacts, poor signal recording, or clinical determination that those electrodes were part of either the SOZ or the IZ. Based on the above, a total of 3731 bipolar contacts were isolated for analysis from the implantations performed in 37 subjects. These bipolar contacts were widely distributed across the lobes with the maximum implant density in temporo-frontal regions. After parceling the brain into 234 brain regions, the minimum number of bipolar contacts in any given parcel were no less than 16.

##### Supplementary figure 5: Electrode distribution

The implantation of invasive electrodes is tailored to the individual patient based on the goal of determining the epileptogenic zone (EZ). This Figure displays the distribution of the invasive electrodes after they are pooled across all subjects (N=37), and grouped according to the lobe in which they are housed.

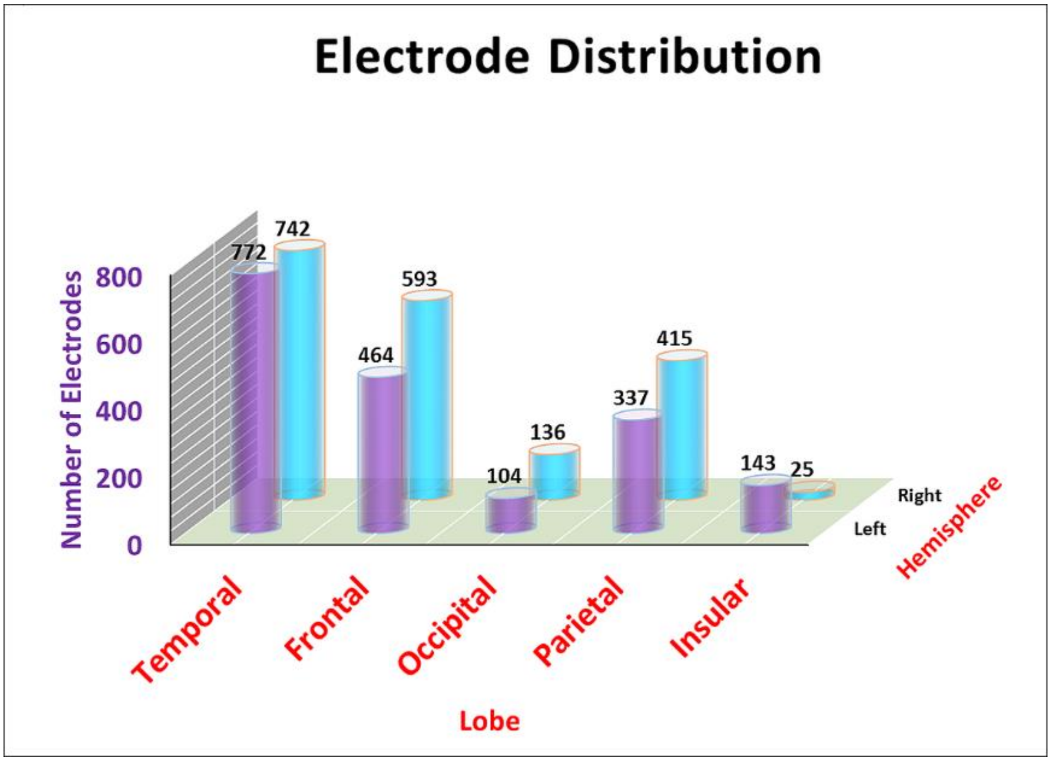

#### 2.3 Understanding hubness of HFA-memory region:

We noted that the HFA-memory network was a composite network involving member nodes from other intrinsic networks. We tried to estimate how the HFA-network compared in its hubness scores to these other networks. The ANOVA model showed the hubness scores (AvgBC) of the different networks in the brain had a distinct range of scores (Supplementary table 2a and b).

**Supplementary Table 2a: AvgBC of HFA-memory network in comparison with the intrinsic networks**

| Multivariate Tests | | | | | | | | | |
|---|---|---|---|---|---|---|---|---|---|
| **Effect** | | **Value** | **F** | **Hypothesis df** | **Error df** | **Sig.** | **Partial Eta Squared** | **Noncent. Parameter** | **Observed Power** |
| Networks | Wilks' Lambda | 0.24 | 23.06 | 9.00 | 65.00 | 0.00 | 0.76 | 207.55 | 1.00 |
| **Tests of Within-Subjects Effects** | | | | | | | | | |
| **Source** | | **Type III Sum of Squares** | **df** | **Mean Square** | **F** | **Sig.** | **Partial Eta Squared** | **Noncent. Parameter** | **Observed Power** |
| Networks | Sphericity Assumed | 512899.13 | 9.00 | 56988.79 | 26.96 | 0.00 | 0.27 | 242.68 | 1.00 |
| | Greenhouse-Geisser | 512899.13 | 6.48 | 79149.01 | 26.96 | 0.00 | 0.27 | 174.74 | 1.00 |
| Error (Networks) | Sphericity Assumed | 1388546.76 | 657.00 | 2113.47 | | | | | |
| | Greenhouse-Geisser | 1388546.76 | 473.05 | 2935.29 | | | | | |
| **Tests of Between-Subjects Effects** | | | | | | | | | |
| **Source** | **Type III Sum of Squares** | **df** | **Mean Square** | **F** | **Sig.** | **Partial Eta Squared** | **Noncent. Parameter** | **Observed Power** | |
| Intercept | 32135516.47 | 1.00 | 32135516.47 | 33397.91 | 0.00 | 1.00 | 33397.91 | 1.00 | |

**Supplementary Table 2b: AvgBC of HFA-memory network in comparison with the other intrinsic networks: post-hoc results**

In this table, we provide the pairwise comparisons of the AvgBC of the different networks..

| Pairwise Comparisons of the BC of the HFA network | | | | | | |
|---|---|---|---|---|---|---|
| Networks | | Mean Difference in AvgBC | Std. Error | Sig. | 95% Confidence Interval for Difference | |
| | | | | | Lower Bound | Upper Bound |
| 1 (TN) | 2 | 2.62 | 10.13 | 0.80 | -17.56 | 22.80 |
| | 3 | -11.18 | 6.85 | 0.11 | -24.83 | 2.46 |
| | 4 | -22.502* | 6.68 | 0.00 | -35.81 | -9.19 |
| | 5 | -27.944* | 7.12 | 0.00 | -42.13 | -13.76 |
| | 6 | -44.223* | 7.86 | 0.00 | -59.89 | -28.56 |
| | 7 | -64.601* | 9.01 | 0.00 | -82.56 | -46.65 |
| | 8 | -62.433* | 7.86 | 0.00 | -78.09 | -46.77 |
| | 9 | -63.984* | 8.31 | 0.00 | -80.54 | -47.43 |
| | 10 | -66.221* | 7.75 | 0.00 | -81.67 | -50.78 |
| 2 (VAN) | 1 | -2.62 | 10.13 | 0.80 | -22.80 | 17.56 |
| | 3 | -13.80 | 8.34 | 0.10 | -30.43 | 2.82 |
| | 4 | -25.122* | 8.32 | 0.00 | -41.71 | -8.53 |
| | 5 | -30.564* | 8.99 | 0.00 | -48.47 | -12.66 |
| | 6 | -46.842* | 9.88 | 0.00 | -66.54 | -27.15 |
| | 7 | -67.220* | 10.39 | 0.00 | -87.92 | -46.52 |
| | 8 | -65.052* | 9.20 | 0.00 | -83.40 | -46.71 |
| | 9 | -66.603* | 10.13 | 0.00 | -86.79 | -46.41 |
| | 10 | -68.840* | 10.03 | 0.00 | -88.82 | -48.86 |
| 3 (VIN) | 1 | 11.18 | 6.85 | 0.11 | -2.46 | 24.83 |
| | 2 | 13.80 | 8.34 | 0.10 | -2.82 | 30.43 |
| | 4 | -11.318* | 5.00 | 0.03 | -21.29 | -1.35 |
| | 5 | -16.760* | 4.91 | 0.00 | -26.54 | -6.98 |
| | 6 | -33.039* | 6.88 | 0.00 | -46.76 | -19.32 |
| | 7 | -53.417* | 7.83 | 0.00 | -69.01 | -37.82 |
| | 8 | -51.249* | 5.54 | 0.00 | -62.28 | -40.21 |
| | 9 | -52.800* | 6.43 | 0.00 | -65.61 | -39.99 |
| | 10 | -55.037* | 5.57 | 0.00 | -66.14 | -43.94 |
| 4 (HFA) | 1 | 22.502* | 6.68 | 0.00 | 9.19 | 35.81 |
| | 2 | 25.122* | 8.32 | 0.00 | 8.53 | 41.71 |

| | | | | | | |
|---|---|---|---|---|---|---|
| | 3 | 11.318* | 5.00 | 0.03 | 1.35 | 21.29 |
| | 5 | -5.44 | 5.22 | 0.30 | -15.84 | 4.95 |
| | 6 | -21.721* | 6.71 | 0.00 | -35.09 | -8.35 |
| | 7 | -42.099* | 7.14 | 0.00 | -56.34 | -27.86 |
| | 8 | -39.930* | 5.34 | 0.00 | -50.57 | -29.29 |
| | 9 | -41.482* | 6.17 | 0.00 | -53.79 | -29.18 |
| | 10 | -43.719* | 5.74 | 0.00 | -55.16 | -32.28 |
| **5 (SMN)** | 1 | 27.944* | 7.12 | 0.00 | 13.76 | 42.13 |
| | 2 | 30.564* | 8.99 | 0.00 | 12.66 | 48.47 |
| | 3 | 16.760* | 4.91 | 0.00 | 6.98 | 26.54 |
| | 4 | 5.44 | 5.22 | 0.30 | -4.95 | 15.84 |
| | 6 | -16.279* | 6.59 | 0.02 | -29.42 | -3.14 |
| | 7 | -36.657* | 7.67 | 0.00 | -51.94 | -21.37 |
| | 8 | -34.488* | 5.89 | 0.00 | -46.24 | -22.74 |
| | 9 | -36.040* | 6.66 | 0.00 | -49.31 | -22.77 |
| | 10 | -38.277* | 6.23 | 0.00 | -50.70 | -25.86 |
| **6 (AN)** | 1 | 44.223* | 7.86 | 0.00 | 28.56 | 59.89 |
| | 2 | 46.842* | 9.88 | 0.00 | 27.15 | 66.54 |
| | 3 | 33.039* | 6.88 | 0.00 | 19.32 | 46.76 |
| | 4 | 21.721* | 6.71 | 0.00 | 8.35 | 35.09 |
| | 5 | 16.279* | 6.59 | 0.02 | 3.14 | 29.42 |
| | 7 | -20.378* | 8.04 | 0.01 | -36.40 | -4.36 |
| | 8 | -18.209* | 7.54 | 0.02 | -33.24 | -3.18 |
| | 9 | -19.761* | 7.29 | 0.01 | -34.30 | -5.22 |
| | 10 | -21.998* | 7.72 | 0.01 | -37.37 | -6.62 |
| **7 (DAN)** | 1 | 64.601* | 9.01 | 0.00 | 46.65 | 82.56 |
| | 2 | 67.220* | 10.39 | 0.00 | 46.52 | 87.92 |
| | 3 | 53.417* | 7.83 | 0.00 | 37.82 | 69.01 |
| | 4 | 42.099* | 7.14 | 0.00 | 27.86 | 56.34 |
| | 5 | 36.657* | 7.67 | 0.00 | 21.37 | 51.94 |
| | 6 | 20.378* | 8.04 | 0.01 | 4.36 | 36.40 |
| | 8 | 2.17 | 8.11 | 0.79 | -14.00 | 18.33 |
| | 9 | 0.62 | 8.50 | 0.94 | -16.32 | 17.56 |
| | 10 | -1.62 | 9.35 | 0.86 | -20.26 | 17.02 |
| **8 (COpN)** | 1 | 62.433* | 7.86 | 0.00 | 46.77 | 78.09 |
| | 2 | 65.052* | 9.20 | 0.00 | 46.71 | 83.40 |
| | 3 | 51.249* | 5.54 | 0.00 | 40.21 | 62.28 |

| | | | | | | |
|---|---|---|---|---|---|---|
| | 4 | 39.930* | 5.34 | 0.00 | 29.29 | 50.57 |
| | 5 | 34.488* | 5.89 | 0.00 | 22.74 | 46.24 |
| | 6 | 18.209* | 7.54 | 0.02 | 3.18 | 33.24 |
| | 7 | -2.17 | 8.11 | 0.79 | -18.33 | 14.00 |
| | 9 | -1.55 | 5.80 | 0.79 | -13.12 | 10.02 |
| | 10 | -3.79 | 6.03 | 0.53 | -15.81 | 8.24 |
| **9 (FPN)** | 1 | 63.984* | 8.31 | 0.00 | 47.43 | 80.54 |
| | 2 | 66.603* | 10.13 | 0.00 | 46.41 | 86.79 |
| | 3 | 52.800* | 6.43 | 0.00 | 39.99 | 65.61 |
| | 4 | 41.482* | 6.17 | 0.00 | 29.18 | 53.79 |
| | 5 | 36.040* | 6.66 | 0.00 | 22.77 | 49.31 |
| | 6 | 19.761* | 7.29 | 0.01 | 5.22 | 34.30 |
| | 7 | -0.62 | 8.50 | 0.94 | -17.56 | 16.32 |
| | 8 | 1.55 | 5.80 | 0.79 | -10.02 | 13.12 |
| | 10 | -2.24 | 6.75 | 0.74 | -15.68 | 11.21 |
| **10 (DMN)** | 1 | 66.221* | 7.75 | 0.00 | 50.78 | 81.67 |
| | 2 | 68.840* | 10.03 | 0.00 | 48.86 | 88.82 |
| | 3 | 55.037* | 5.57 | 0.00 | 43.94 | 66.14 |
| | 4 | 43.719* | 5.74 | 0.00 | 32.28 | 55.16 |
| | 5 | 38.277* | 6.23 | 0.00 | 25.86 | 50.70 |
| | 6 | 21.998* | 7.72 | 0.01 | 6.62 | 37.37 |
| | 7 | 1.62 | 9.35 | 0.86 | -17.02 | 20.26 |
| | 8 | 3.79 | 6.03 | 0.53 | -8.24 | 15.81 |
| | 9 | 2.24 | 6.75 | 0.74 | -11.21 | 15.68 |

**Supplementary Table 3: Betweenness Centrality of the HFA-memory regions compared to the rest of the intrinsic networks**

After determining the functional composition of the HFA-memory network we sought to find how the nodes in the HFA-memory network behaved differently from the network they originate from in terms of their Betweenness Centrality. We found that FPN and COpN nodes that shared membership with the HFA-memory network had lower BC compared to the non-member nodes of the respective networks while the member nodes of TN had higher BC compared to the non-member nodes.

| | **T** | **p** |
|---|---|---|
| HFA_TN vs Other_TN | +2.85 | 0.043 |
| HFA_VIN vs Other_VIN | -0.29 | 0.77 |
| HFA_SMN vs Other_SMN | 1.04 | 0.20 |
| HFA_DAN vs Other_DAN | -1.14 | 0.14 |
| ***HFA_COpN vs Other_COpN**** | -3.47 | $5.2*10^{-4}$ |
| ***HFA_FPN vs Other_FPN**** | -3.63 | $2.9*10^{-4}$ |
| HFA_DMN vs Other_DMN | -1.1 | 0.25 |

T – T-score from T-test; p= probability value; HFA_TN- nodes of temporal network that are members of HFA-memory network; HFA_VIN- nodes of visual integrated network that are members of HFA-memory network; HFA_SMN- nodes of somatomotor network that are members of HFA-memory network; HFA_DAN- nodes of dorsal attention network that are members of HFA-memory network; HFA_COpN- nodes of cingulo-opercular network that are members of HFA-memory network; HFA_FPN- nodes of frontoparietal network that are members of HFA-memory network; HFA_DMN- nodes of default mode network that are members of HFA-memory network

**Supplementary figure 6: Two-Way Factorial analysis**

The BC levels of HFA-members of intrinsic networks (HFA_COpN and HFA_FPN) differed from the remaining regions of a given intrinsic network (Other_COpN, Other_FPN). However, what was more important to note was, if this effect was contributed by the PAT vs CON group difference. Hence we tested a Two-Way Factorial analysis. We found that The HFA-memory regions still differed in terms of the regions but there was no significant difference across the groups as noted in both Figure 6a and Figure 6b. In Figure 6a, we note that the main effect of BC of HFA_COpN was lower than Others_COpN (F=12.56, p=0.001) while the main effect of PAT vs CON difference was not significant (F=1.33, p=0.25) nor the interaction was significant (F=0.15, p=0.69). Similarly, in Figure 6b, we note that the main effect of BC of HFA_FPN was lower than Others_FPN (F=19.53, $p=1.9*10^{-5}$) while the main effect of PAT vs CON difference was not significant (F=0.27, p=0.65) nor the interaction was significant (F=0.09, p=0.76).

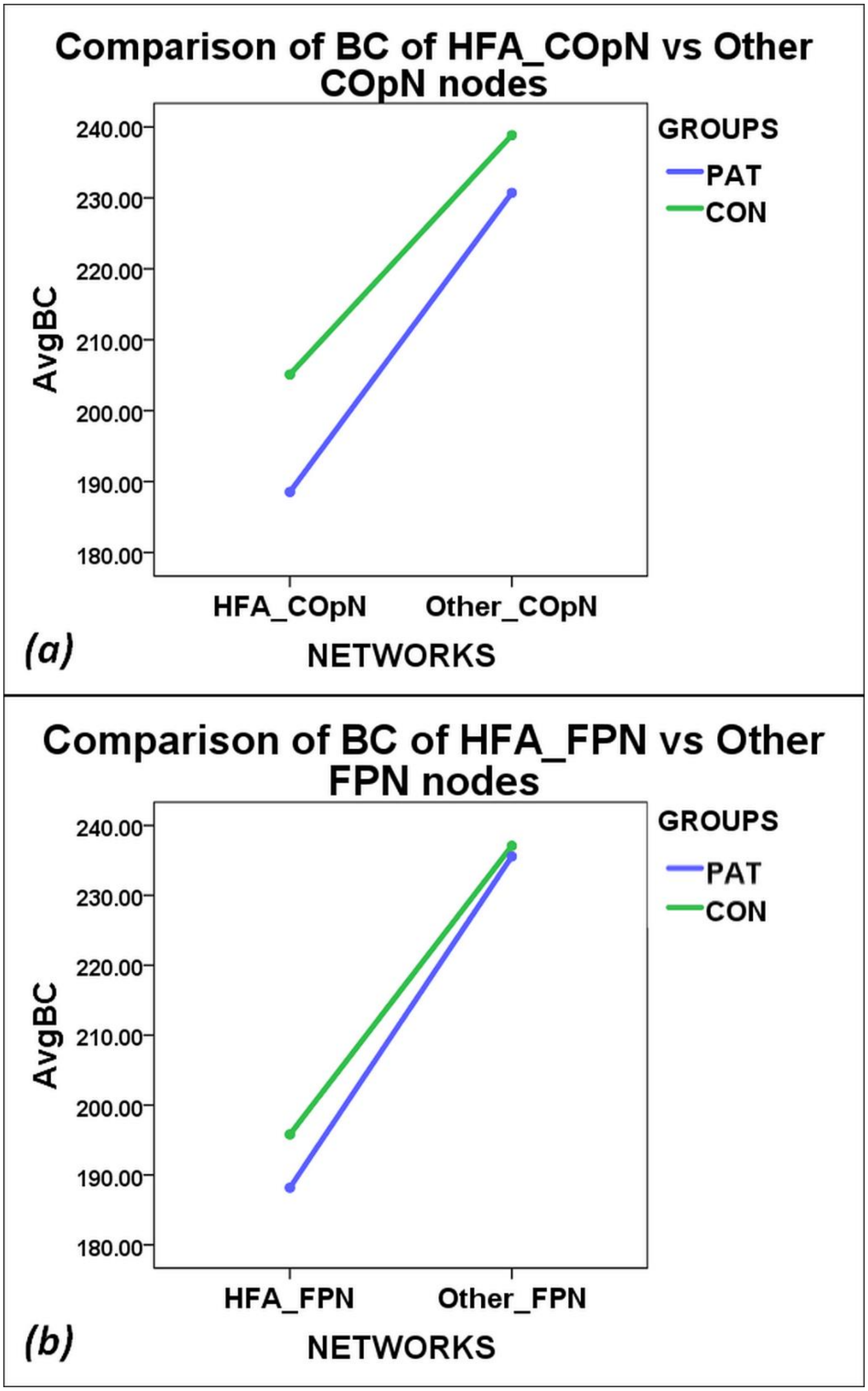
Comparison of BC of HFA_COpN vs Other COpN nodes
AvgBC
240.00
230.00
220.00
210.00
200.00
190.00
180.00
GROUPS
PAT
CON
HFA_COpN
Other_COpN
NETWORKS
(a)
Comparison of BC of HFA_FPN vs Other FPN nodes
AvgBC
240.00
230.00
220.00
210.00
200.00
190.00
180.00
GROUPS
PAT
CON
HFA_FPN
Other_FPN
NETWORKS
(b)

**Supplementary Table 3a: Average Participation Coefficient of HFA-memory network in comparison with the other intrinsic networks**

| Multivariate Tests | | | | | | | | | |
|---|---|---|---|---|---|---|---|---|---|
| Effect | | Value | F | Hypothesis df | Error df | Sig. | Partial Eta Squared | Noncent. Parameter | Observed Power |
| Networks | Wilks' Lambda | 0.14 | 44.87 | 9.00 | 65.00 | 0.00 | 0.86 | 403.84 | 1.00 |
| **Tests of Within-Subjects Effects** | | | | | | | | | |
| **Source** | | **Type III Sum of Squares** | **df** | **Mean Square** | **F** | **Sig.** | **Partial Eta Squared** | **Noncent. Parameter** | **Observed Power** |
| Networks | Sphericity Assumed | 0.11 | 9.00 | 0.01 | 50.47 | 0.00 | 0.41 | 454.25 | 1.00 |
| | Greenhouse-Geisser | 0.11 | 5.14 | 0.02 | 50.47 | 0.00 | 0.41 | 259.53 | 1.00 |
| Error(Networks) | Sphericity Assumed | 0.16 | 657.00 | 0.00 | | | | | |
| | Greenhouse-Geisser | 0.16 | 375.37 | 0.00 | | | | | |

| Tests of Between-Subjects Effects | | | | | | | | |
|---|---|---|---|---|---|---|---|---|
| Source | Type III Sum of Squares | df | Mean Square | F | Sig. | Partial Eta Squared | Noncent. Parameter | Observed Power |
| Intercept | 501.96 | 1.00 | 501.96 | 408028.28 | 0.00 | 1.00 | 408028.28 | 1.00 |
| Error | 0.09 | 73.00 | 0.00 | | | | | |

**Supplementary Table 3b: Average Participation Coefficient of HFA-memory network in comparison with the other intrinsic networks: post-hoc results**

In this table, we provide the pairwise comparisons of the Average Participation Coefficient (AvgPC) of the different networks.

| Pairwise Comparisons of the AvgPC of the HFA network | | | | | | |
|---|---|---|---|---|---|---|
| Networks | | Mean Difference in Average Participation Coefficient | Std. Error | Sig. | 95% Confidence Interval for Difference | |
| | | | | | Lower Bound | Upper Bound |
| 1 (TN) | 2 | 0.013* | 0.00 | 0.00 | 0.01 | 0.02 |
| | 3 | 0.025* | 0.00 | 0.00 | 0.02 | 0.03 |
| | 4 | 0.001 | 0.00 | 0.30 | 0.00 | 0.00 |
| | 5 | 0.013* | 0.00 | 0.00 | 0.01 | 0.02 |
| | 6 | 0.015* | 0.00 | 0.00 | 0.01 | 0.02 |
| | 7 | -0.016* | 0.00 | 0.00 | -0.02 | -0.01 |
| | 8 | -0.011* | 0.00 | 0.00 | -0.02 | -0.01 |
| | 9 | -0.007* | 0.00 | 0.00 | -0.01 | 0.00 |
| | 10 | -0.001 | 0.00 | 0.81 | -0.01 | 0.00 |
| 2 (VAN) | 1 | -0.013* | 0.00 | 0.00 | -0.02 | -0.01 |
| | 3 | 0.012* | 0.00 | 0.00 | 0.01 | 0.02 |
| | 4 | -0.011* | 0.00 | 0.00 | -0.02 | -0.01 |
| | 5 | 0.000 | 0.00 | 0.90 | -0.01 | 0.01 |
| | 6 | 0.002 | 0.00 | 0.61 | -0.01 | 0.01 |
| | 7 | -0.029* | 0.00 | 0.00 | -0.04 | -0.02 |
| | 8 | -0.024* | 0.00 | 0.00 | -0.03 | -0.02 |
| | 9 | -0.020* | 0.00 | 0.00 | -0.03 | -0.01 |
| | 10 | -0.014* | 0.00 | 0.00 | -0.02 | -0.01 |
| 3 (VIN) | 1 | -0.025* | 0.00 | 0.00 | -0.03 | -0.02 |
| | 2 | -0.012* | 0.00 | 0.00 | -0.02 | -0.01 |
| | 4 | -0.024* | 0.00 | 0.00 | -0.03 | -0.02 |
| | 5 | -0.012* | 0.00 | 0.00 | -0.02 | -0.01 |
| | 6 | -0.010* | 0.00 | 0.00 | -0.02 | 0.00 |

| | | | | | | |
|---|---|---|---|---|---|---|
| | 7 | -0.041* | 0.00 | 0.00 | -0.05 | -0.04 |
| | 8 | -0.036* | 0.00 | 0.00 | -0.04 | -0.03 |
| | 9 | -0.032* | 0.00 | 0.00 | -0.04 | -0.03 |
| | 10 | -0.026* | 0.00 | 0.00 | -0.03 | -0.02 |
| **4 (HFA)** | 1 | -0.001 | 0.00 | 0.30 | 0.00 | 0.00 |
| | 2 | 0.011* | 0.00 | 0.00 | 0.01 | 0.02 |
| | 3 | 0.024* | 0.00 | 0.00 | 0.02 | 0.03 |
| | 5 | 0.012* | 0.00 | 0.00 | 0.01 | 0.02 |
| | 6 | 0.014* | 0.00 | 0.00 | 0.01 | 0.02 |
| | 7 | -0.018* | 0.00 | 0.00 | -0.02 | -0.02 |
| | 8 | -0.013* | 0.00 | 0.00 | -0.02 | -0.01 |
| | 9 | -0.009* | 0.00 | 0.00 | -0.01 | -0.01 |
| | 10 | -0.002 | 0.00 | 0.38 | -0.01 | 0.00 |
| **5 (SMN)** | 1 | -0.013* | 0.00 | 0.00 | -0.02 | -0.01 |
| | 2 | 0.000 | 0.00 | 0.90 | -0.01 | 0.01 |
| | 3 | 0.012* | 0.00 | 0.00 | 0.01 | 0.02 |
| | 4 | -0.012* | 0.00 | 0.00 | -0.02 | -0.01 |
| | 6 | 0.002 | 0.00 | 0.52 | 0.00 | 0.01 |
| | 7 | -0.030* | 0.00 | 0.00 | -0.03 | -0.03 |
| | 8 | -0.024* | 0.00 | 0.00 | -0.03 | -0.02 |
| | 9 | -0.021* | 0.00 | 0.00 | -0.03 | -0.02 |
| | 10 | -0.014* | 0.00 | 0.00 | -0.02 | -0.01 |
| **6 (AN)** | 1 | -0.015* | 0.00 | 0.00 | -0.02 | -0.01 |
| | 2 | -0.002 | 0.00 | 0.61 | -0.01 | 0.01 |
| | 3 | 0.010* | 0.00 | 0.00 | 0.00 | 0.02 |
| | 4 | -0.014* | 0.00 | 0.00 | -0.02 | -0.01 |
| | 5 | -0.002 | 0.00 | 0.52 | -0.01 | 0.00 |
| | 7 | -0.031* | 0.00 | 0.00 | -0.04 | -0.03 |
| | 8 | -0.026* | 0.00 | 0.00 | -0.03 | -0.02 |
| | 9 | -0.023* | 0.00 | 0.00 | -0.03 | -0.02 |
| | 10 | -0.016* | 0.00 | 0.00 | -0.02 | -0.01 |
| **7 (DAN)** | 1 | 0.016* | 0.00 | 0.00 | 0.01 | 0.02 |
| | 2 | 0.029* | 0.00 | 0.00 | 0.02 | 0.04 |
| | 3 | 0.041* | 0.00 | 0.00 | 0.04 | 0.05 |
| | 4 | 0.018* | 0.00 | 0.00 | 0.02 | 0.02 |

| | | | | | | |
|---|---|---|---|---|---|---|
| | 5 | 0.030* | 0.00 | 0.00 | 0.03 | 0.03 |
| | 6 | 0.031* | 0.00 | 0.00 | 0.03 | 0.04 |
| | 8 | 0.005* | 0.00 | 0.00 | 0.00 | 0.01 |
| | 9 | 0.009* | 0.00 | 0.00 | 0.01 | 0.01 |
| | 10 | 0.016* | 0.00 | 0.00 | 0.01 | 0.02 |
| **8 (COpN)** | 1 | 0.011* | 0.00 | 0.00 | 0.01 | 0.02 |
| | 2 | 0.024* | 0.00 | 0.00 | 0.02 | 0.03 |
| | 3 | 0.036* | 0.00 | 0.00 | 0.03 | 0.04 |
| | 4 | 0.013* | 0.00 | 0.00 | 0.01 | 0.02 |
| | 5 | 0.024* | 0.00 | 0.00 | 0.02 | 0.03 |
| | 6 | 0.026* | 0.00 | 0.00 | 0.02 | 0.03 |
| | 7 | -0.005* | 0.00 | 0.00 | -0.01 | 0.00 |
| | 9 | 0.004* | 0.00 | 0.01 | 0.00 | 0.01 |
| | 10 | 0.010* | 0.00 | 0.00 | 0.01 | 0.01 |
| **9 (FPN)** | 1 | 0.007* | 0.00 | 0.00 | 0.00 | 0.01 |
| | 2 | 0.020* | 0.00 | 0.00 | 0.01 | 0.03 |
| | 3 | 0.032* | 0.00 | 0.00 | 0.03 | 0.04 |
| | 4 | 0.009* | 0.00 | 0.00 | 0.01 | 0.01 |
| | 5 | 0.021* | 0.00 | 0.00 | 0.02 | 0.03 |
| | 6 | 0.023* | 0.00 | 0.00 | 0.02 | 0.03 |
| | 7 | -0.009* | 0.00 | 0.00 | -0.01 | -0.01 |
| | 8 | -0.004* | 0.00 | 0.01 | -0.01 | 0.00 |
| | 10 | 0.007* | 0.00 | 0.00 | 0.00 | 0.01 |
| **10 (DMN)** | 1 | 0.001 | 0.00 | 0.81 | 0.00 | 0.01 |
| | 2 | 0.014* | 0.00 | 0.00 | 0.01 | 0.02 |
| | 3 | 0.026* | 0.00 | 0.00 | 0.02 | 0.03 |
| | 4 | 0.002 | 0.00 | 0.38 | 0.00 | 0.01 |
| | 5 | 0.014* | 0.00 | 0.00 | 0.01 | 0.02 |
| | 6 | 0.016* | 0.00 | 0.00 | 0.01 | 0.02 |
| | 7 | -0.016* | 0.00 | 0.00 | -0.02 | -0.01 |
| | 8 | -0.010* | 0.00 | 0.00 | -0.01 | -0.01 |
| | 9 | -0.007* | 0.00 | 0.00 | -0.01 | 0.00 |

**Supplementary figure 7: AvgPC score of the different networks in an ascending order**

This box and whisker plot shows the ranking of the different network on the basis of their AvgPC. Cognitive control networks such as FPN, COpN, and DAN have a higher AvgPC compared to sensorimotor processing networks such as SMN, AN, and VIN. The HFA-memory network has an AvgPC value that ranks 5th and is similar to that of TN and DMN (marked by the yellow highlighted box).

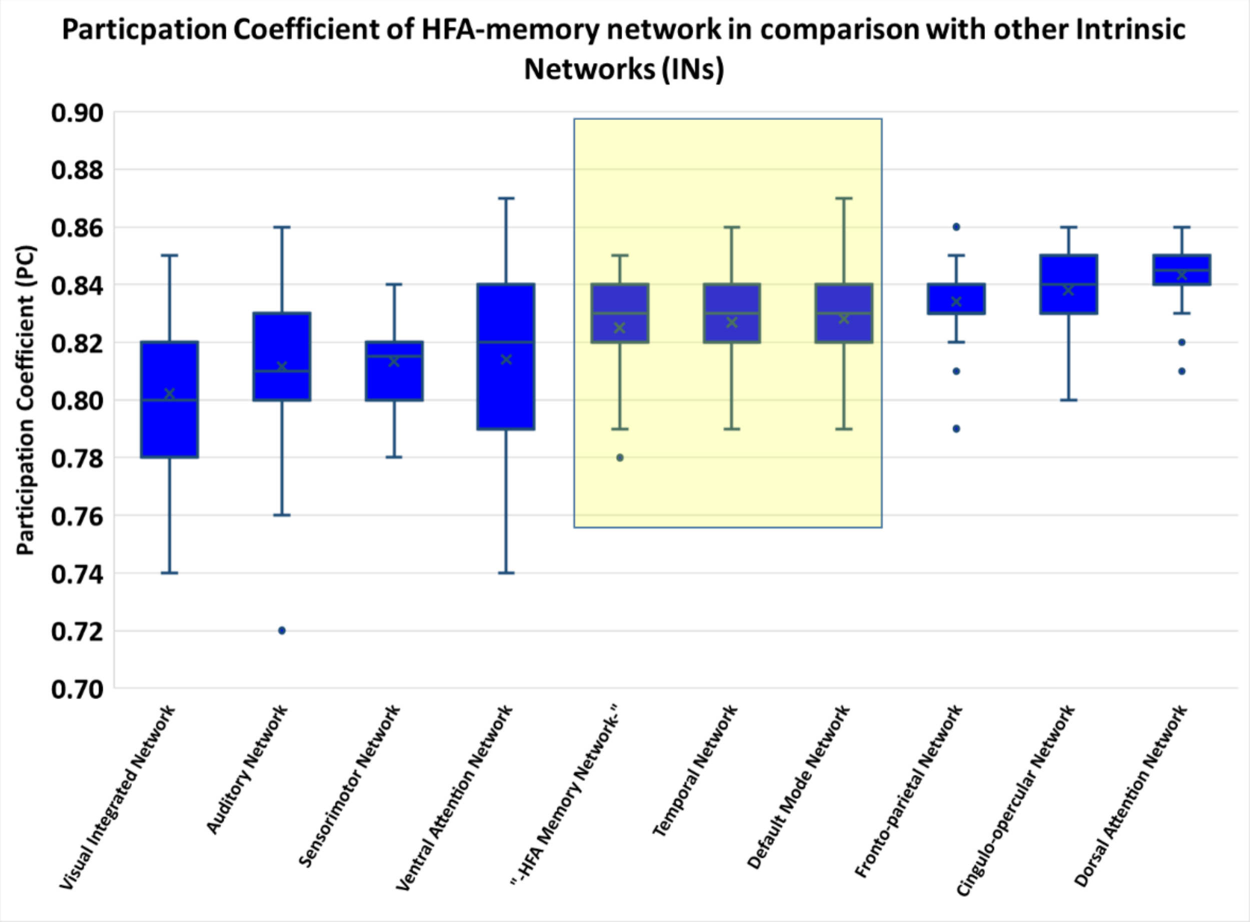

#### Supplementary figure 8: Memory encoding network from a meta-analysis of task fMRI

The memory encoding network from a meta-analysis 124 studies on task-based fMRI for memory encoding is presented here (results are corrected using FDR (FDR <0.01)(Yarkoni T et al., 2011). The regions involved are BL_Amygdala, BL_Angular, BL_Frontal_Inf_Oper, BL_Frontal_Inf_Orb, BL_Frontal_Inf_Tri, BL_Fusiform, BL_Hippocampus, BL_Insula, BL_Occipital_Mid, BL_Occipital_Sup, BL_ParaHippocampal, BL_Parietal_Inf, BL_Parietal_Sup, BL_Precentral, BL_Supp_Motor_Area, L_Caudate, L_Cerebelum, L_Frontal_Sup_Medial, L_Lingual, L_Occipital_Inf, L_Pallidum and L_Putament. In our study, the hippocampus was not significant for SME because it was actually found that in 11/37 subjects hippocampal electrodes were excluded from analysis as they a part of the ictogenic area. Hence, the hippocampal electrodes were excluded and the net SME of hippocampus was not statistically significant.

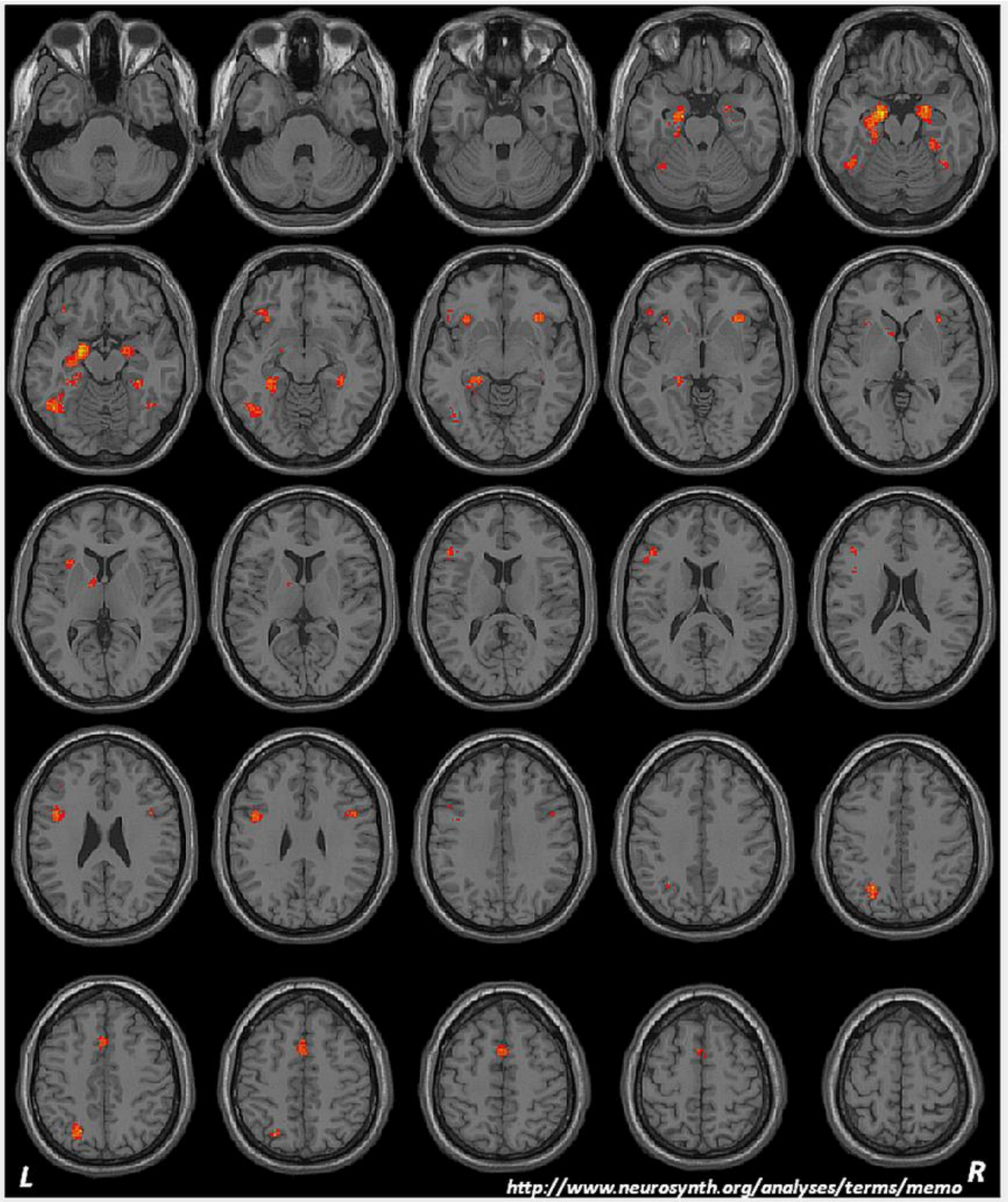
L
R
http://www.neurosynth.org/analyses/terms/memo

##### 2.3.1 References:

Alexander-Bloch A, Giedd JN, Bullmore E (2013), Imaging structural co-variance between human brain regions. Nat Rev Neurosci 14:322-336.

Alexander-Bloch AF, Gogtay N, Meunier D, Birn R, Clasen L, Lalonde F, Lenroot R, Giedd J, et al. (2010), Disrupted modularity and local connectivity of brain functional networks in childhood-onset schizophrenia. Frontiers in systems neuroscience 4:147.

AlexanderBloch AF, Gogtay N, Meunier D, Birn R, Clasen L, Lalonde F, Lenroot R, Giedd J, et al. (2010), Disrupted modularity and local connectivity of brain functional networks in childhood-onset schizophrenia. Frontiers in systems neuroscience 4:147.

Altmann A, Tolosi L, Sander O, Lengauer T (2010), Permutation importance: a corrected feature importance measure. Bioinformatics 26:1340-1347.

Ashburner J, Friston KJ (2005), Unified segmentation. Neuroimage 26:839-851.

Burke JF, Ramayya AG, Kahana MJ (2015), Human intracranial high-frequency activity during memory processing: neural oscillations or stochastic volatility? Curr Opin Neurobiol 31:104-110.

Friston KJ, Williams S, Howard R, Frackowiak RS, Turner R (1996), Movement-related effects in fMRI time-series. Magnetic resonance in medicine 35:346-355.

Hagmann P, Cammoun L, Gigandet X, Meuli R, Honey CJ, Wedeen VJ, Sporns O (2008), Mapping the structural core of human cerebral cortex. PLoS biology 6:e159.

He X, Doucet GE, Pustina D, Sperling MR, Sharan AD, Tracy JI (2017), Presurgical thalamic “hubness” predicts surgical outcome in temporal lobe epilepsy. Neurology 88:2285-2293.

Kragel JE, Ezzyat Y, Sperling MR, Gorniak R, Worrell GA, Berry BM, Inman C, Lin JJ, et al. (2017), Similar patterns of neural activity predict memory function during encoding and retrieval. Neuroimage 155:60-71.

Rondinoni C, Amaro E, Jr., Cendes F, dos Santos AC, Salmon CE (2013), Effect of scanner acoustic background noise on strict resting-state fMRI. Braz J Med Biol Res 46:359-367.

Rubinov M, Sporns O (2010), Complex network measures of brain connectivity: uses and interpretations. Neuroimage 52:1059-1069.

Stam C, Tewarie P, Van Dellen E, Van Straaten E, Hillebrand A, Van Mieghem P (2014), The trees and the forest: characterization of complex brain networks with minimum spanning trees. International Journal of Psychophysiology 92:129-138.

Su L, Wang L, Shen H, Feng G, Hu D (2013), Discriminative analysis of non-linear brain connectivity in schizophrenia: an fMRI Study. Front Hum Neurosci 7:702.

Tewarie P, van Dellen E, Hillebrand A, Stam CJ (2015), The minimum spanning tree: an unbiased method for brain network analysis. Neuroimage 104:177-188.

Tolosi L, Lengauer T (2011), Classification with correlated features: unreliability of feature ranking and solutions. Bioinformatics 27:1986-1994.

van Diessen E, Diederen SJ, Braun KP, Jansen FE, Stam CJ (2013), Functional and structural brain networks in epilepsy: what have we learned? Epilepsia 54:1855-1865.

Yan CG, Cheung B, Kelly C, Colcombe S, Craddock RC, Di Martino A, Li Q, Zuo XN, et al. (2013), A comprehensive assessment of regional variation in the impact of head micromovements on functional connectomics. Neuroimage 76:183-201.

Yan CG, Zang YF (2010), DPARSF: A MATLAB Toolbox for "Pipeline" Data Analysis of Resting-State fMRI. Frontiers in systems neuroscience 4:13.

Yarkoni T, Poldrack RA, Nichols TE, Van Essen DC, Wager TD (2011), Large-scale automated synthesis of human functional neuroimaging data. Nat Methods 8:665-670.
